# Mitotic Block and Epigenetic Repression Underlie Neurodevelopmental Defects and Neurobehavioral Deficits in Congenital Heart Disease

**DOI:** 10.1101/2023.11.05.565716

**Authors:** George C. Gabriel, Hisato Yagi, Tuantuan Tan, Abha S. Bais, Benjamin J. Glennon, Margaret C. Stapleton, Lihua Huang, William T. Reynolds, Marla G. Shaffer, Madhavi Ganapathiraju, Dennis Simon, Ashok Panigrahy, Yijen L. Wu, Cecilia W. Lo

## Abstract

Poor neurodevelopment is often observed with congenital heart disease (CHD), especially with mutations in chromatin modifiers. Here analysis of mice with hypoplastic left heart syndrome (HLHS) arising from mutations in Sin3A associated chromatin modifier *Sap130*, and adhesion protein *Pcdha9,* revealed neurodevelopmental and neurobehavioral deficits reminiscent of those in HLHS patients. Microcephaly was associated with impaired cortical neurogenesis, mitotic block, and increased apoptosis. Transcriptional profiling indicated dysregulated neurogenesis by REST, altered CREB signaling regulating memory and synaptic plasticity, and impaired neurovascular coupling modulating cerebral blood flow. Many neurodevelopmental/neurobehavioral disease pathways were recovered, including autism and cognitive impairment. These same pathways emerged from genome-wide DNA methylation and Sap130 chromatin immunoprecipitation sequencing analyses, suggesting epigenetic perturbation. Mice with *Pcdha9* mutation or forebrain-specific *Sap130* deletion without CHD showed learning/memory deficits and autism-like behavior. These novel findings provide mechanistic insights indicating the adverse neurodevelopment in HLHS may involve cell autonomous/nonautonomous defects and epigenetic dysregulation and suggest new avenues for therapy.

## Introduction

Congenital heart disease (CHD) affects up to 1% of live births^1^. With recent surgical advances that allow infants born with CHD to survive to adulthood, there are now more adults living with CHD than infants born annually with CHD. Unfortunately, CHD survivors often suffer developmental delay with high risk of neurodevelopmental impairment that can significantly degrade quality of life^2^. Thus, a “neurobehavioral signature” has been described that includes learning disabilities, impaired social/communication skills, autism spectrum disorders, and other cognitive, behavioral, and neuropsychiatric deficits^2,3^. Lack of mechanistic insights into the causes for the neurodevelopmental impairment has precluded the development of targeted therapy for early intervention^2^.

The major contributing factors for the poor neurodevelopment were previously thought to arise from hypoxic injury and complications related to multiple cardiac surgeries. However, outcome studies have pointed to patient intrinsic factors^3-6^. This is consistent with the findings of small head circumference, microcephaly, and brain abnormalities identified in utero before cardiac surgery^7-10^. That these patient intrinsic factors may encompass genetic factors is suggested by the finding of *de novo* pathogenic variants significantly associated with CHD, such as in chromatin modifiers^11,12^. Interestingly, chromatin modifiers also are reported to cause autism, intellectual disability, cognitive impairment, and neuropsychiatric disorders outside of the CHD population^11-13^. In addition, genetic disorders and other patient-specific factors such as socioeconomic status have been reported to increase the risk for adverse neurodevelopmental outcome in CHD^6,14^.

Among patients with CHD, the highest morbidity and mortality is observed with hypoplastic left heart syndrome (HLHS), a severe CHD now survivable with a three-stage surgical palliation. However, HLHS survivors have high risk of neurodevelopmental delay and neurobehavioral and cognitive impairment^2^. This is associated with reductions in functional status and quality of life, increase in behavioral symptoms, poor adaptive skills required for navigating school and daily living, and intellectual disability with deficits in full scale IQ^15-17^. HLHS is also associated with impaired brain development. Microcephaly, holoprosencephaly, and agenesis of the corpus callosum are observed in 25% of HLHS fetuses^2,10^. Microcephaly was shown to predict early adverse neurologic outcomes^18^. While the brain volume reductions were correlated with lower cerebral substrate delivery, neurodevelopmental outcome was not improved after in utero fetal intervention with aortic valvuloplasty, suggesting intrinsic brain deficits^19^.

To investigate the potential causes of poor neurodevelopment in HLHS, here we leverage the *Ohia* mouse model of HLHS. This mouse model was recovered from a large scale forward genetic screen for CHD and was shown to harbor two recessive mutations that in combination cause HLHS, Sin3A-associated chromatin modifier *Sap130,* and protocadherin cell adhesion protein *Pcdha9* from the α-protocadherin gene cluster^20^. Both the Sin3A complex and the clustered protocadherins have essential roles in development of the nervous system. The Sin3A protein is part of the histone deacetylase (HDAC) repressor complex and is essential for brain development. Thus, *SIN3A* deficiency causes Witteveen-Kolk syndrome, a neurodevelopmental disorder associated with intellectual disability, autism, microcephaly, and facial dysmorphism^21^. Also found in the Sin3A complex is MECP2, an X-linked methyl CpG binding protein associated with Rett syndrome, the most common cause of cognitive impairment in females^22^. Further, the Sin3A complex is known to regulate DNAm via interactions with the Tet family of methylcytosine dioxygenases, and DNA methylation (DNAm) signatures have been shown to be diagnostic of neurodevelopmental/neuropsychiatric disorders^23-25^. The clustered protocadherins also play critical roles in development of the nervous system. They are cell adhesion proteins encoding cell surface diversity that specify neuronal identity patterning synaptic connectivity in the brain^26^. Mice with deletion of the entire *Pcdha* gene cluster exhibit deficits in synaptic connectivity and behavioral impairment^27^. Interestingly, *PCDHA* and *SIN3A* mutations are clinically implicated in autism and in CHD^21,28-30^.

Leveraging the *Ohia* mouse model, we examined the impact of *Sap130/Pcdha9* mutations on development of the brain, examining structure and differentiation of the brain with histopathological analyses, immunoconfocal microscopy, and molecular profiling with RNA sequencing (RNAseq), Sap130 chromatin immunoprecipitation sequencing (ChIPseq), and genome wide DNA methylation analyses. To investigate the role of *Sap130* in the microcephaly phenotype, we generated mice with forebrain-specific deletion of *Sap130*, and as the latter mice and mice harboring only the *Ohia Pcdha9* mutation are adult viable, functional effects of the *Sap130* and *Pcdha9* mutations were examined with neurobehavioral assessments. Together these studies yielded novel insights into the cellular and molecular mechanisms contributing to the poor neurodevelopmental outcome clinically associated with HLHS.

## Results

*Ohia* mutant mice double homozygous for the *Sap130/Pcdha9* mutations (*Ohia^m/m^)* were previously shown to exhibit HLHS and other CHD with incomplete penetrance^20^. Analysis of double homozygous mutants (n=83) showed 56.7% displayed craniofacial/head defects. Of these, 48.2% exhibited both CHD and craniofacial/head defects, 8.5% showed only craniofacial/head defects, and 24% showed only CHD. Severity of the craniofacial/head defects was variable, ranging from mild micrognathia to severe agnathia often accompanied by low set ears, dome shaped head, and eye defects (Figure S1). Examination of gross brain anatomy in *Ohia* mice (n=40) from E14.5 to E18.5 revealed 81% have forebrain hypoplasia, and 59% have severe microcephaly (Figure 1A-D). In half of the severe cases (49.6%) holoprosencephaly was also observed. Analysis of severe mutants using episcopic confocal microscopy (ECM) for serial section histological 3D reconstruction revealed hypoplastic/aplastic olfactory bulbs, thin cerebral cortex, hypoplasia of the cerebellum, reduction of cerebellar fissures, dilation of the lateral ventricles, and hypoplasia of the corpus callosum (Figure 1E-J). Quantification confirmed significant cortical thinning, indicating defects in cortical plate development (Figure 1K-N).

**Figure 1.**
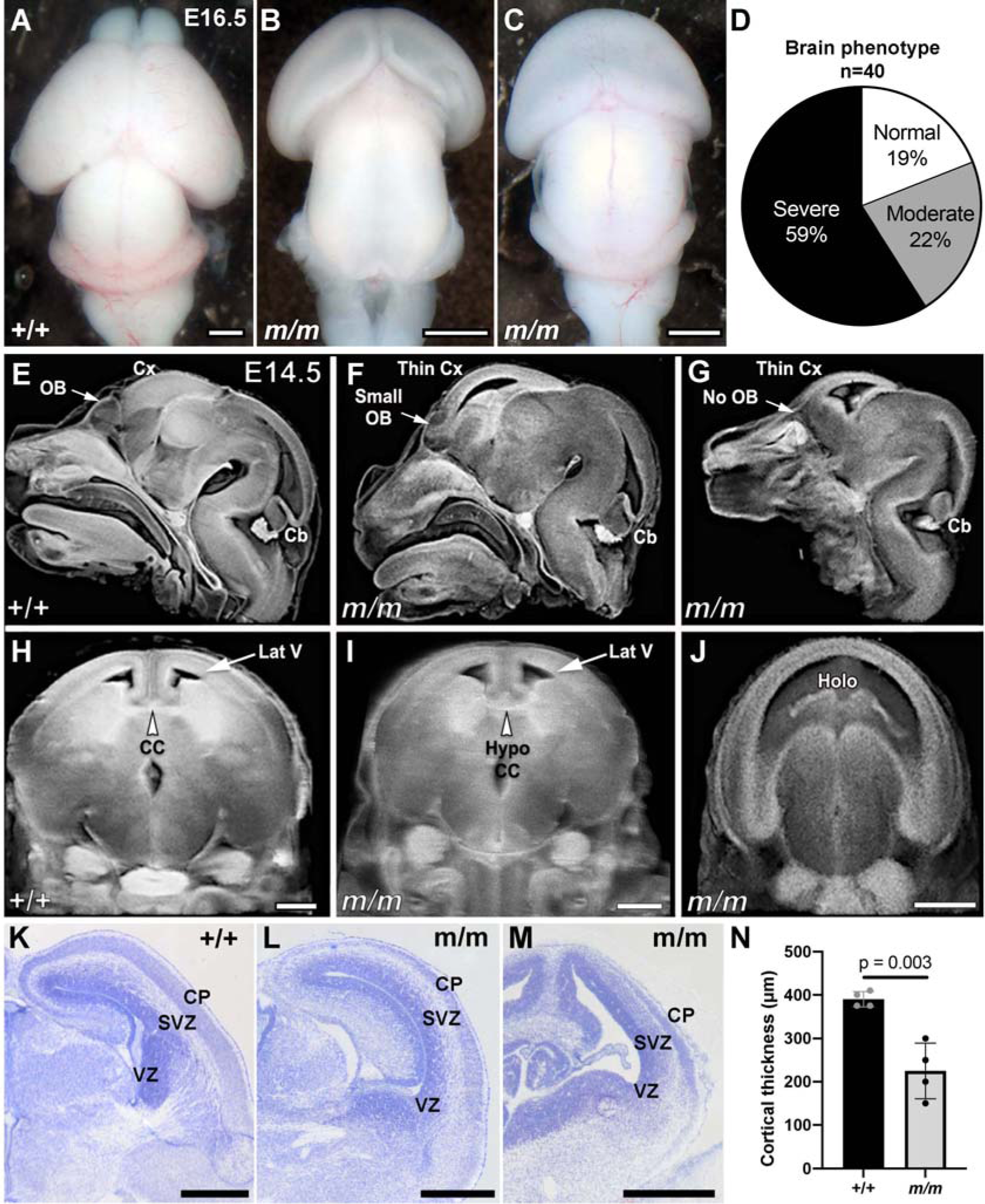
*Ohia^m/m^* mice display microcephaly with brain abnormalities. **A-D.** E16.5 wildtype (A) and *Ohia^m/m^* mutant mouse brains exhibiting moderate (B) and severe (C) forebrain hypoplasia, with the severe mutant also showing holoprosencephaly. The distribution of brain phenotype severity in *Ohia^m/m^* mice is shown in the pie chart (D). **E-J.** Episcopic confocal imaging of the head shown in the sagittal and coronal plane of a wildtype (+/+) (E, H) and two *Ohia^m/m^* mice, one with mild (F, I) and the other with severe (G, J) forebrain hypoplasia. Note hypoplastic (F) or absent (G) olfactory bulb, thin cortex (F,G), hypoplastic corpus callosum (I), and holoprosencephaly (J). **K-N.** Cresyl violet stained sections from control (K), and two *Ohia* mutants with mild (L) vs. severe (M) forebrain hypoplasia. Quantification of 5 *Ohia* mutants and 5 littermate controls showed reduced cortical thickness (N). p=0.003, 2-tailed unpaired t-test. Cx, cortex; OB, olfactory bulb; Cb, cerebellum; CC, corpus callosum; Lat V, lateral ventricle; Holo, holoprosencephaly. Scale bars = 0.5mm.

### Impaired Cortical Neurogenesis with Reduction of Intermediate Progenitors

Cortical plate formation is orchestrated via the expansion of neural progenitors in the ventricular and subventricular zone of the developing brain. These progenitors give rise to cells that migrate upwards to form layers II through VI of the cortical plate, with the innermost layer, layer VI, being the first to form, and Layer II, the outermost layer being the last to form. Immunostaining using antibodies for different progenitor lineages and different layers of the cortical plate showed no change in the Pax6+ apical progenitors (radial glial cells) in the ventricular zone, but Tbr2+ intermediate progenitors in the subventricular zone were markedly reduced (Figure 2A-L). Satb2, a marker for postmitotic neurons in the upper layers of the developing cortex was also markedly reduced, indicating failure to form the later born neurons in layers II-IV (Fig 2Q-T vs. M-P, quantified in Fig 2U-Y). In contrast, the Tbr1+ earlier born neurons of layers VI, and Ctip2+ neurons in layer V-VI were present, but abnormally distributed in overlapping domains as compared to their normal pattern of deployment in wildtype mice (Figure 2Q-T vs. M-P). These observations indicate a cortical neurogenesis defect in the *Ohia^m/m^* mutant brain involving failure of the intermediate progenitors to expand, thereby causing deficiency in the later born neurons in layers II-IV.

**Figure 2.**
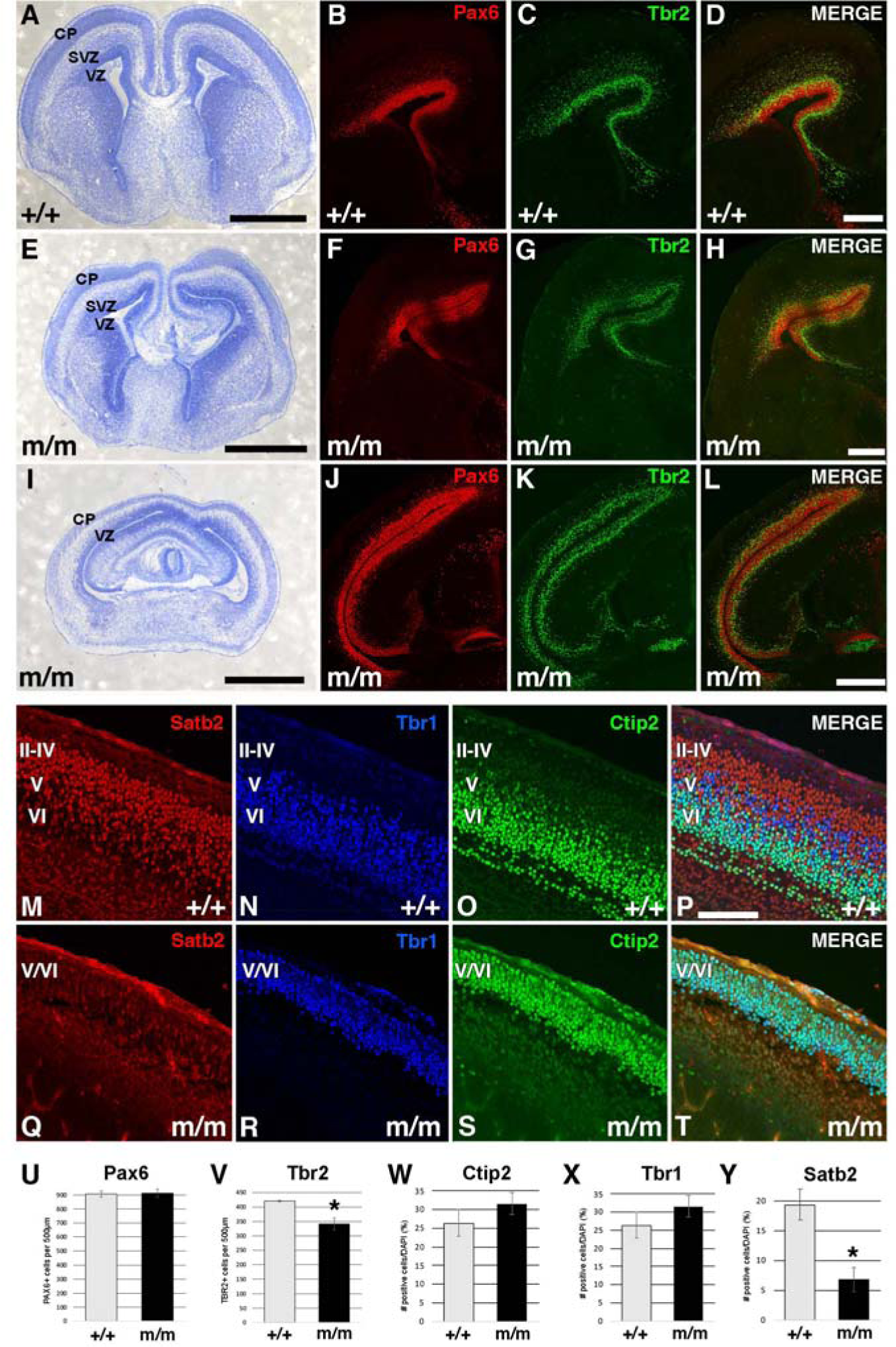
Impaired Cortical Plate Formation with Loss of Tbr2+ Intermediate Progenitors in *Ohia^m/m^*Mutant Brain. **A-L**. Sections of the brain of E16.5 control (A-D) and *Ohia^m/m^* mutant mice with mild (E-H) or severe (I-L) forebrain hypoplasia were stained with cresyl violet (A,E,I) to delineate the brain tissue architecture. Antibody staining was conducted for markers of apical progenitor cells (Pax6, B,F,J), intermediate progenitor cells (Tbr2 in C,G,K), and merged image showing both markers (D,H,L). **M-T.** Section of the cortical plate of E16.5 wildtype (M-P) and mutant (Q-T) mouse brain was stained with antibodies to Satb2, marker for cortical layers II-IV (M,Q), Ctip2 for cortical layers V-VI (N,R), and Tbr1 (O,S) for cortical layer VI, and merged image showing all three markers (P,T). **U-Y.** Quantification of number of cells stained by Pax6 (U), Tbr2 (V), Ctip2 (W), Tbr1 (X), and Satb2 (Y) in the brain of 5 *Ohia^m/m^* mutant and 5 littermate controls. P-values calculated by 2-tailed, unpaired t-test. Scale bars in A-L indicate 500µm. Scale bars in M-T indicate 100µm.

### Loss of Neural Progenitors Associated with Mitotic Block

To elucidate the mechanism of the cortical neurogenesis defect in the *Ohia^m/m^* brain, we quantified cell proliferation and apoptosis. Analysis of E14.5 *Ohia^m/m^* brain tissue showed a significant increase in pH3 positive cells in the subventricular zone, and also increased TUNEL throughout the ventricular and subventricular zones (Figure 3A-F, quantified in G), findings reminiscent of those seen in the *Ohia^m/m^*HLHS heart tissue^20^. The fraction of mitotic cells in anaphase/telophase decreased, indicating metaphase block (Figure 3H)^20^. Similar findings have been reported in mouse models of primary microcephaly and are observed to be associated with mitotic spindle defects, such as with mutations in *ASPM*, Assembly Factor for Spindle Microtubules, and *WDR62*, another centrosome/spindle protein, two genes that account for over 50% of primary microcephaly^31,32^.

**Figure 3.**
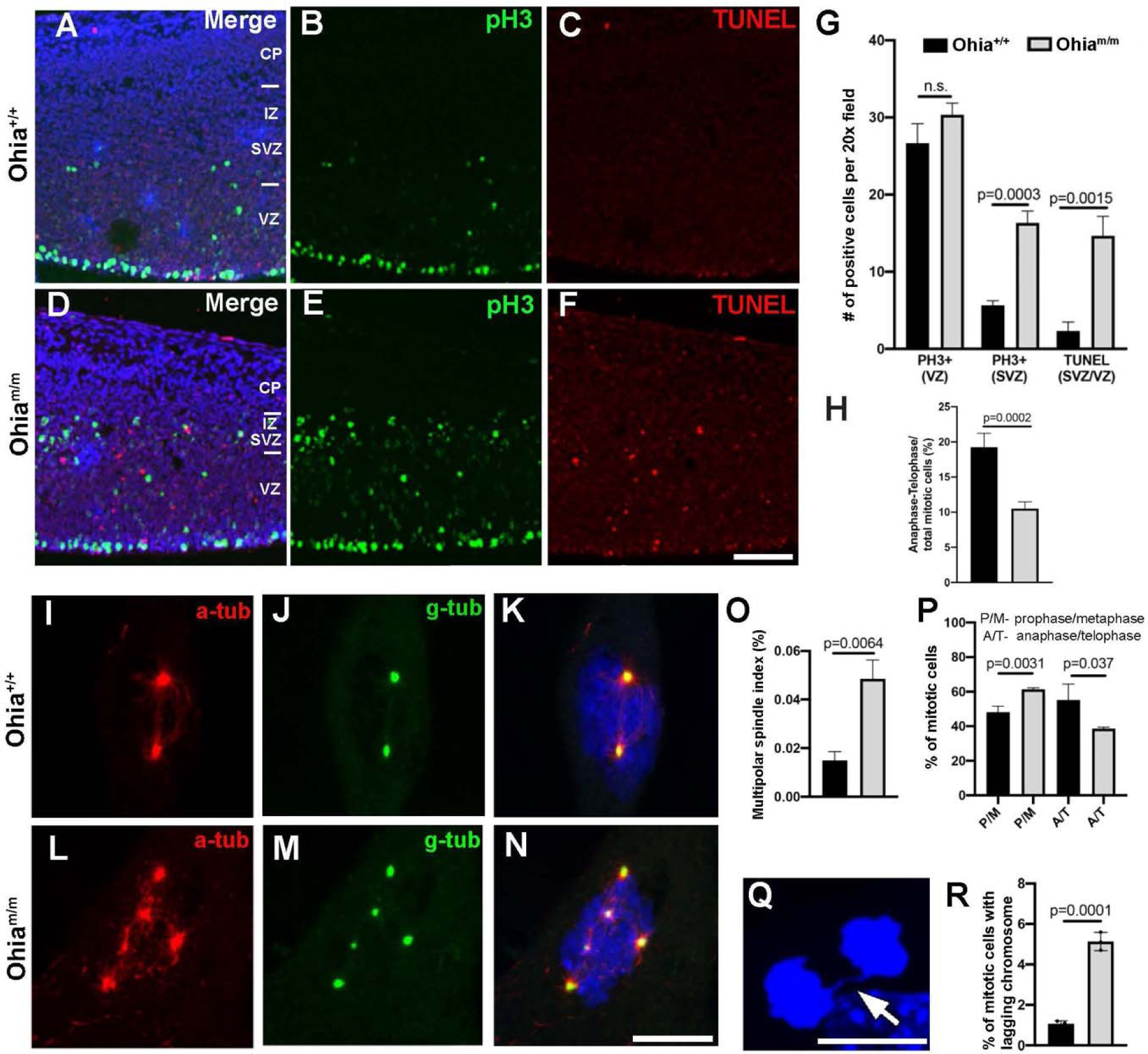
Neural Progenitors in the Cortical Plate Show Mitotic Block With Increase in Cell Death and Possible Role for Multipolar Spindles Indicated by MEF Analysis. **A-H.** Comparison of the cortex of the brain from E14.5 *Ohia^m/m^* mutant (D-F) vs. wildtype littermate control embryos (A-C) showed more pH3 and TUNEL staining (G) in the ventricular (VZ) and subventricular (SVC) zones in the mutants. A decrease was also noted in the proportion of mitotic cells in anaphase-telophase (H). p-value obtained using 2-way, unpaired t-test. (n=3, *Ohia^m/m^*mutants, n=3 littermate controls). **I-N.** MEFs generated from wildtype (I-K) and *Ohia^m/m^* mutant embryos (L-N) were immunostained with α-tubulin and γ-tubulin. Mitotic cells in the mutant MEFs showed increase in multipolar spindles (quantification in panel O). MEFs were generated from 3 *Ohia^m/m^* and 3 littermate control embryos, with >500 cells analyzed per embryo derived MEF. P-value calculated by 2-way, unpaired t-test. **P-R.** *Ohia^m/m^* MEFs showed reduction in the proportion of mitotic cells in anaphase/telophase (P). and an increase in lagging chromosomes during anaphase (Q,R) P-value obtained using 2-way, unpaired t-test. CP, cortical plate; IZ, intermediate zone; SVZ, subventricular zone; VZ, ventricular zone; P/M, prophase/metaphase; A/T, anaphase/telophase. Scale bars = 100µm.

To investigate for spindle defects, we generated mouse embryonic fibroblasts (MEFs) derived from *Ohia^m/m^* mutant embryos and littermate controls. The MEFs generated were immunostained for α-tubulin to visualize the spindle apparatus and γ-tubulin for the centrosome. This revealed a marked increase in multi-polar spindles in the *Ohia^m/m^* MEFs, which is seldom seen in wildtype MEFs (Figure 3L-N vs. 3I-K, quantified in 3O). This was associated with an increase of mitotic cells at prophase/metaphase (P/M), while mitotic cells at anaphase/telophase (A/T) decreased, indicating a mitotic block (Figure 3P). A mitotic spindle defect was indicated by the further observation of a significant increase in lagging chromosomes in mitotic cells at anaphase (Figure 3Q, R). Together these findings support a spindle defect contributing to the loss of later born neurons in cortical layers II-IV underlying the emergence of microcephaly in *Ohia^m/m^* mutant mice.

### Dysregulated Gene Expression and Defects in Neurodevelopment

Transcriptome profiling was conducted with RNAseq analysis of E13.5-E14.5 brain tissue from *Ohia^m/m^*mutants with severe brain phenotype (n=3) and wildtype littermate controls (n=5). This yielded 1,549 differentially expressed genes (DEGs), 1,081 downregulated and 468 upregulated (FDR 0.05; Supplemental Spreadsheet 1). Metascape analysis of the DEGs yielded many nervous system related terms, including forebrain development, forebrain generation of neurons, synapse assembly, axon guidance, action potential, glutamatergic synapses, ion transport, and learning and memory (Figure 4A). Ingenuity Pathway Analysis (IPA) recovered as upstream regulators, REST - repressor element 1 silencing transcription factor, which was upregulated, and DTNBP1 - dystrobrevin binding protein 1, and SGK1 - serum glucocorticoid regulated kinase 1, both downregulated (Figure 4B). The recovery of REST in the transcriptome profiling is particularly notable in the context of microcephaly. REST controls expansion of neural progenitors via recruitment of HDACs to the Sin3A complex^33,34^. This would predict the transcriptional repression of proneuronal genes, including the GRIN family of glutamate receptors observed to be downregulated in the *Ohia* mutant brain (Figure 4B)^35^. The upstream regulator DTNBP1 regulates neurotransmitter release and receptor signaling and is linked to cognitive impairment and schizophrenia, while SGK1 regulates glutamate receptor expression and facilitates learning and hippocampal long-term potentiation^36,37^. Together, these upstream regulators and their downstream genes are predicted to perturb biological processes that are all neurobehavioral related, including learning, long-term potentiation, coordination, conditioning, analgesia, hyperalgesia (Figure 4B).

**Figure 4.**
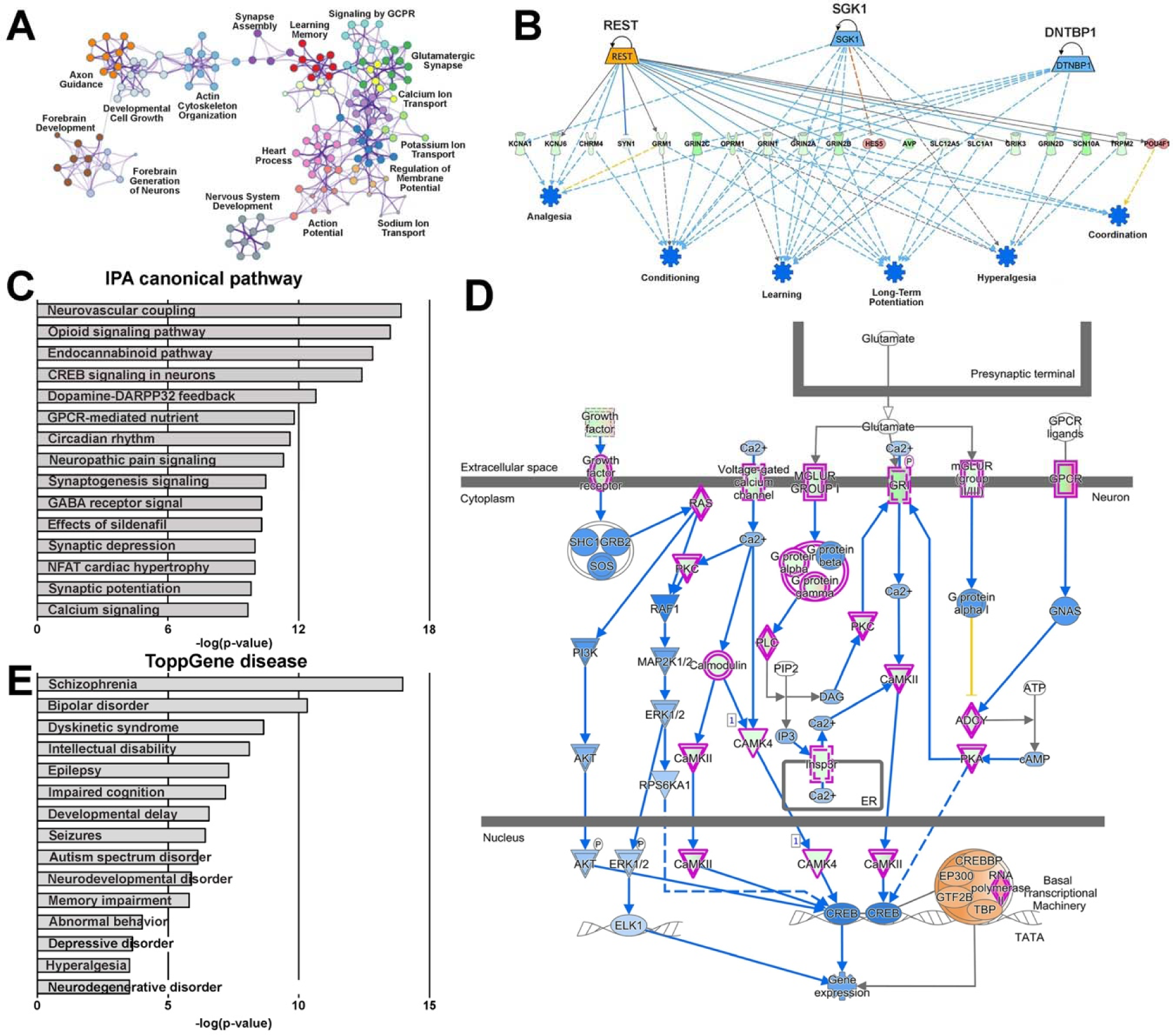
Transcriptome Profiling Shows Dysregulated Neurodevelopment in *Ohia^m/m^* Mutants. Differentially expressed genes (DEGs) recovered from RNASeq analysis of E14.5 *Ohia* mutant brain were analyzed for pathway enrichment using Metascape (A), Ingenuity Pathway Analysis (IPA) for upstream regulators of downstream neurodevelopmental outcomes (B, red for upregulated, green for downregulated), IPA canonical pathway (C), IPA curated CREB signaling pathway (D, purple outline indicates genes found to be differentially expressed, green fill indicates downregulated and red fill indicates upregulated DEGs), as well as ToppGene Disease analysis (E).

Further canonical pathway analysis using IPA recovered cyclic AMP response element binding protein (CREB) signaling, synaptic depression/potentiation, synaptogenesis signaling, GABA receptor signaling, and calcium signaling (Figure 4C; Supplemental Dataset 1). Associated with the CREB signaling pathway is reduction in glutamate receptor signaling and downregulation of PKC, PLC, PKA, calmodulin, and CAMKII/CAMK4 (Figure 4D). This pathway has critical roles in synaptic plasticity mediating long term memory, and is involved in the pathogenic alteration of synaptic plasticity and memory associated with neurocognitive disorders and various neurodegenerative diseases such as Alzheimer’s Disease and Huntington’s disease, and also implicated in schizophrenia and depressive disorders^38^. Interestingly, this analysis also recovered circadian rhythm, likely a reflection of the known role of CREB in circadian rhythm regulation (Figure 4C)^39^. ToppGene disease pathway enrichment analysis also identified autism, mental depression, intellectual disability, seizures, neurodegenerative diseases, and neuropsychiatric diseases such as schizophrenia and bipolar disorder (Figure 4E, Supplemental Dataset 1).

### NO Signaling Defect and the Perturbation of Neurovascular Coupling

The top IPA pathway recovered from the *Ohia* mutant brain RNAseq analysis was neurovascular coupling, an autoregulatory mechanism by which the brain’s energy demands are closely coordinated with cerebral blood flow (Figure 4C,5, Supplemental Figure S2)^40^. Such cerebral autoregulation is mediated by neuronal activity dependent production of nitric oxide (NO), which promotes vascular smooth muscle relaxation and dilation of cerebral arteries to increase blood flow to the brain. This cerebral autoregulation, which has been described in patients with some types of CHD, results in enhanced cerebral perfusion providing a brain sparing effect^41^. In the *Ohia* mutant brain, we observed marked downregulation of NO synthase 1 (NOS1) (Figure 5; FDR=8.03E-14, Supplemental Spreadsheet 1) and guanylate cyclase, a NO receptor and sensor generating cGMP, the second messenger mediating NO signaling with downstream protein kinase G (PKG) activation to modulate vasodilation and neuronal activity (Figure 5). In contrast to NOS and guanylate cyclase, PKG expression was upregulated, perhaps a compensatory response to the reduction in NO and cGMP (Figure 5). IPA pathway analysis also recovered the “Effect of sildenafil”, a drug commonly used to promote vasodilation by enhancing NO levels via inhibition of phosphodiesterase breaking down cGMP (Figure 4C). We note NO signaling has additional roles in CREB regulated neuronal gene expression important in learning/memory, and thus NO deficiency is also expected to cause learning/memory deficits^42^.

**Figure 5.**
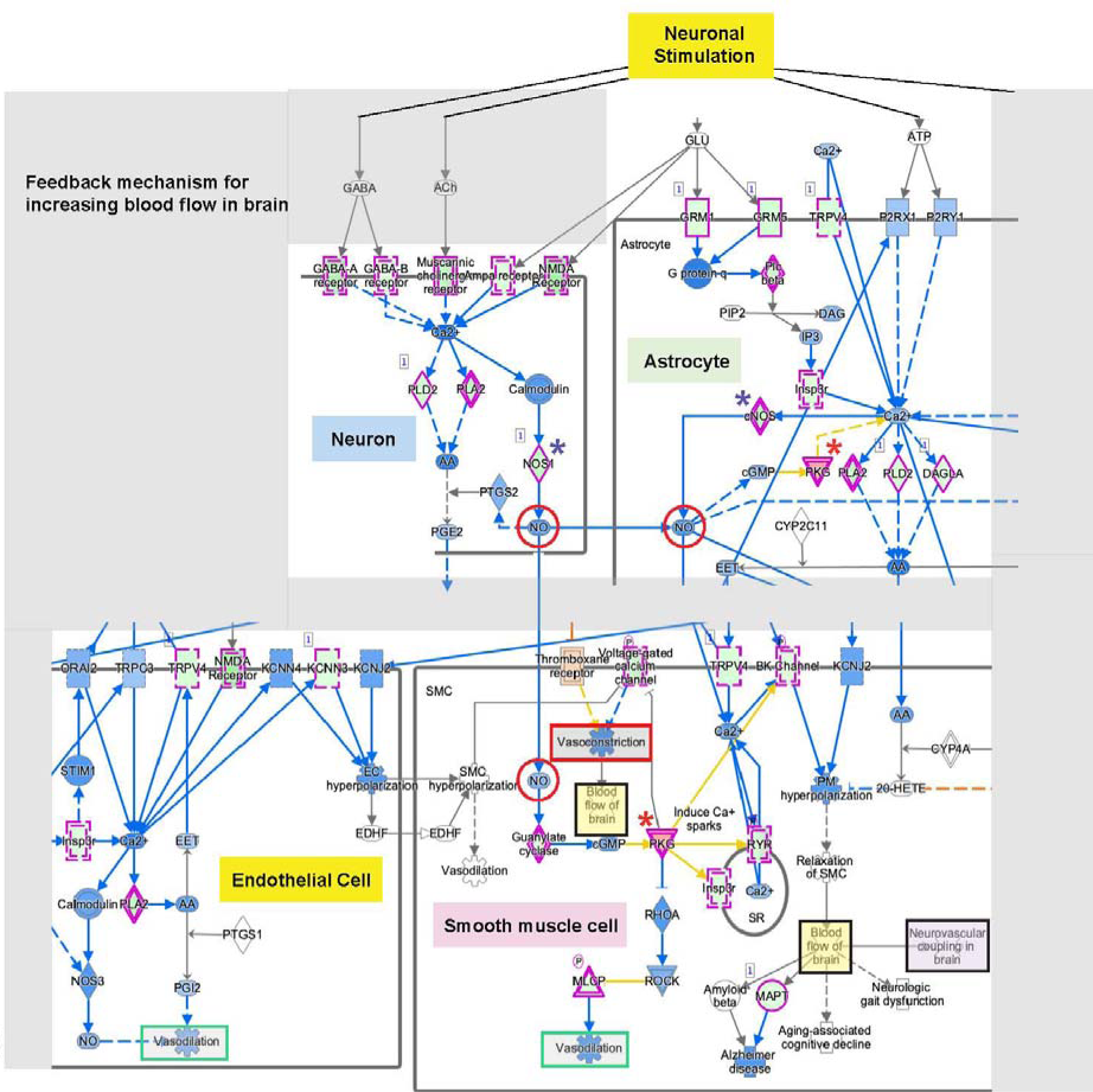
Dysregulated Expression of Genes Associated with Neurovascular Coupling. Modified IPA curated neurovascular coupling pathway illustrating dysregulated expression of genes involved in neurovascular coupling. DEGs identified in *Ohia* brain highlighted in purple outline.

### Correlation of DNA Methylation with Transcriptomic Profiles and Brain Phenotypes

We investigated for possible involvement of epigenetic regulation in the brain defects of the *Ohia* mutant mice, as the Sin3A-HDAC repressor complex to which Sap130 is bound is known to regulate DNAm with binding of Tet DNA demethylases. This was investigated with methylome analysis of DNA from the forebrain tissue of E15.5 *Ohia^m/m^* mutant mice (n=3) with severe microcephaly phenotype vs. C57BL/6J wildtype mice (n=3) using the Illumina Infinium mouse methylome array containing probes for over 285,000 CpG sites genome wide^43^. This analysis recovered 5,117 differentially methylated regions (DMRs) associated with 4,179 genes (FDR<0.05; Supplemental Spreadsheet 2). By way of example, we show CpG methylation associated with a DMR in a highly conserved genomic region at the terminal exon of *Klf13*, a known negative regulator of CREB signaling essential for axonogenesis and neurite outgrowth (Figure 6A)^44^.

**Figure 6.**
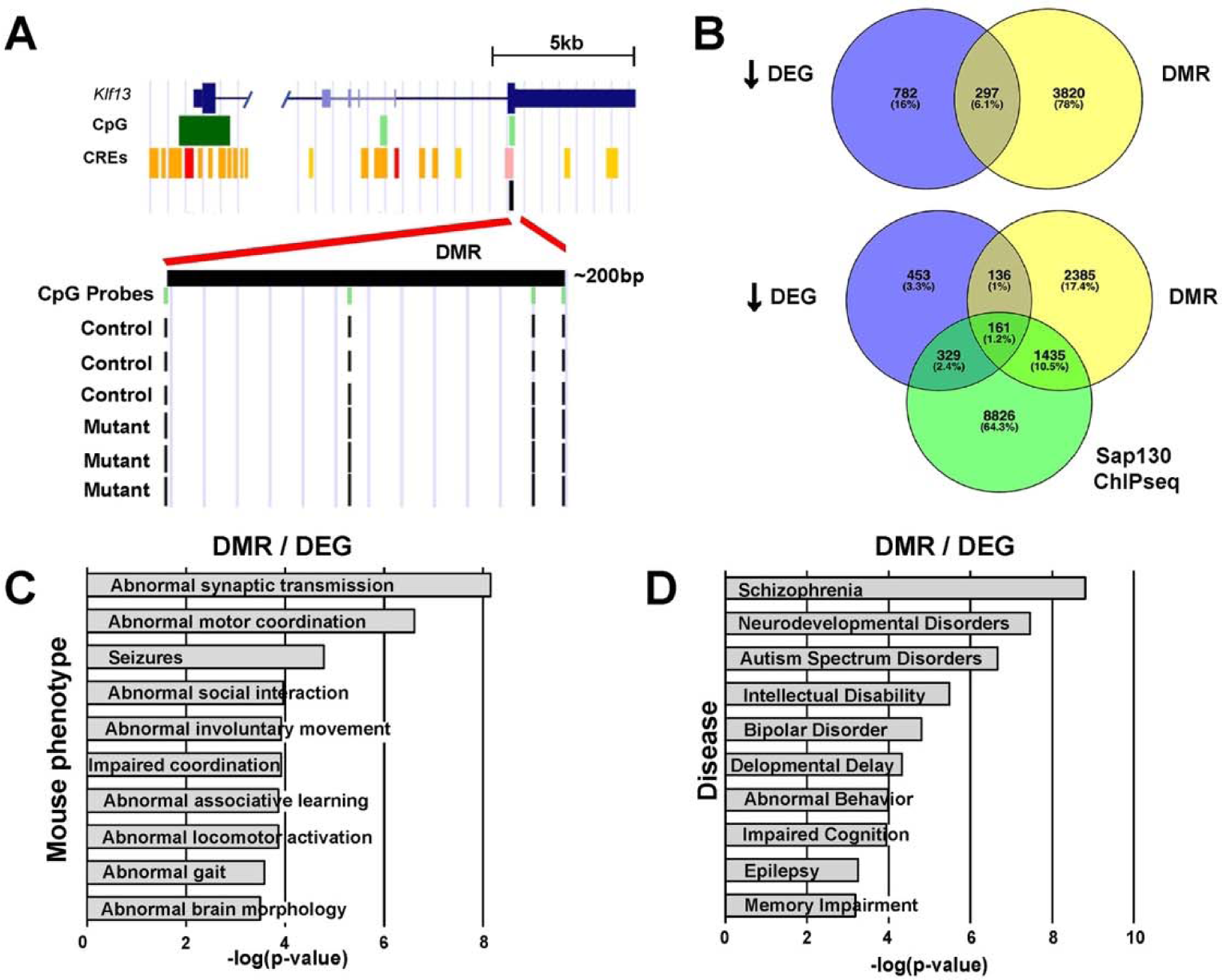
Molecular profiling of *Ohia^m/m^* brain tissue with integration of genome-wide methylome with RNAseq and Sap130 ChIPseq analyses. **A.** Increased methylation of CpG sites in a DMR found in *Klf17* in the *Ohia^m/m^* brain tissue. **B.** Venn diagrams show two-way intersection of downregulated DEGs with genes associated with DMR and three-way intersection of down regulated DEGs, genes associated with DMR, and genes recovered from Sap130 ChiPSeq. **C, D.**ToppGene enrichment analysis for Mouse Phenotypes (C) and Diseases (D) recovered from the two-way intersection of the downregulated DEGs with genes associated with DMRs.

Intersection of DEGs that are downregulated with DMR associated genes identified 297 genes (Figure 6B, Supplemental Spreadsheet 3). ToppGene pathway enrichment analysis of these 297 intersecting genes for Mouse Phenotypes recovered terms related to brain, behavior, and neurological function (Figure 6C, Supplemental Spreadsheet 3), and for Disease, we observed autism, intellectual disability, epilepsy, and other neurological diseases known to be associated with HLHS (Figure 6D; Supplemental Spreadsheet 3). In contrast, only 61 genes were recovered from interrogating the DEG/DMR intersecting genes comprising the upregulated DEGs, with pathway enrichment analysis recovering mostly metabolic pathways (Figure S3, Supplemental Spreadsheet 3).

These observations suggest DNAm changes associated with the downregulated genes may drive the adverse neurodevelopmental outcome associated with HLHS. To investigate this possibility further, we analyzed DNAm in *Ohia* mutants with mild phenotype comprising those with little or no forebrain hypoplasia (Supplemental Figure S4). DMR analysis yielded only 1,921 genes as compared to 4,179 genes for DNAm analysis of the severe brain phenotype mutants (Figure S5A). In the severe mutants, 2,539 (60%) of the 4,179 genes were exclusive to the severe phenotype, but for the mild brain phenotype, only 343 (17.9%) of 1,921 genes were exclusive to the mild brain phenotype. Analysis of all the DMR associated genes in the mild phenotype mutants using ToppGene DisGeNET focused analysis yielded only 6 of 10 top neurological diseases obtain from all DMR associated genes of the severe phenotype, all showing much lower p values (Supplemental Spreadsheet 4). For example, the p value for schizophrenia is e-23 for severe and e-9 for mild, or for neurodevelopmental disorder, e-07 for severe and e-02 for mild (Supplemental Spreadsheet 4). Similar interrogation of DMRs exclusive to the severe or mild mutants yielded no significant disease terms for mild, while the same disease terms with lower p values were observed for the severe phenotype (Figure S5B; Supplemental Spreadsheet 4). Additional analysis of DMR associated genes shared between the mild and severe mutants yielded only two disease terms, schizophrenia and intellectual disability, both with low p values (Figure S5B; Supplemental Spreadsheet 4). Overall, these observations support DNAm playing a role in the transcriptional changes associated with *Ohia* mutant brain phenotype severity.

### Sap130 Epigenetic Regulation of Neurogenesis and Neurological Disease Pathways

We further investigated whether Sap130 may contribute to the epigenetic regulation by conducting Sap130 chromatin immunoprecipitation sequencing (ChIPseq) analysis to identify potential targets of Sap130 regulation. Sap130 ChIPseq analysis of E13.5 wildtype mouse brain tissue yielded 16,231 Sap130 binding sites encompassing 10,753 genes (Supplemental Spreadsheet 5). Among the genes recovered were transcription regulators of neurogenesis such as *Rest*, *Sin3A*, *CTCF, Satb2, Klf13,* and also microcephaly-related genes such as *Aspm* and *Wdr62* (Figure 7A). Transcription factor binding site analysis identified enrichment for SP, KLF, IRF, ELF, and RFX motifs (Figure 7B). SP and KLF are closely related zinc finger proteins that bind GC rich promoters, and are commonly associated with pluripotency and cell cycle genes^45^. KLFs, Kruppel like transcription factors, also have important roles in brain and heart development, with KLF2 regulating vascular tone in response to blood flow^44,46,47^.

**Figure 7.**
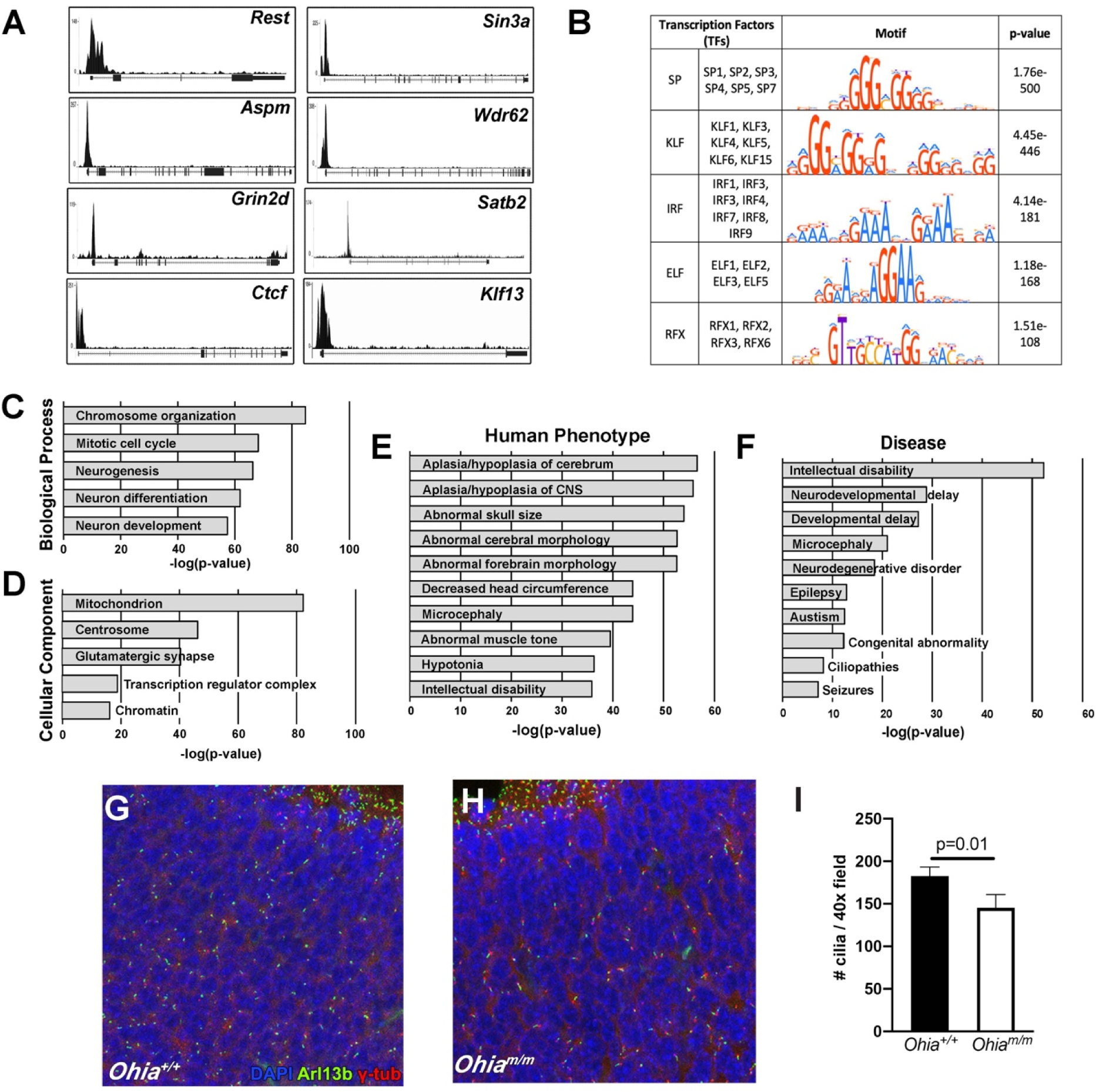
Sap130 Chromatin Immunoprecipitation Sequencing Analysis of Mouse Brain and Examination of Cilia in the *Ohia* Brain Tissue. **A.** Sap130 occupancy in the 5’ upstream promoter region of selected genes is shown in panel A. **B.** Motif enrichment analysis of the Sap130 ChIPseq target genes identified putative Sap130 DNA binding sequences. **C-F.** Pathways in GO Biological Process (C) and Cellular Components (D), as well as Human Phenotype (E), and Disease (F) terms recovered from ToppGene gene enrichment analysis of Sap130 ChIPseq target genes. **G-I.** Cilia visualization in sections of wildtype (G) and *Ohia^m/m^* mutant brain tissue (H) immunostained with Arl13b (cilia, green), γ-tub (centrosome, red) and DAPI. Quantification showed reduction of cilia in the mutant brain (I, n=3 *Ohia* mutant and 4 littermate controls). P value determined by 2-tailed, unpaired t-test.

Pathway enrichment analysis yielded GO biological processes comprising chromosome organization, cell cycle progression, and neurogenesis (Figure 7C). Cellular components identified included mitochondrion, centrosome, glutamatergic synapse and transcriptional regulation (Figure 7D). Human Phenotype and Disease Pathway enrichment analyses identified many of the same terms seen with the down regulated DEGs, such as microcephaly and intellectual disability (Figure 7E). Among Human Phenotype recovered were aplasia/hypoplasia of the cerebrum, decreased head circumference, hypotonia, and for Disease terms, neurodevelopmental delay, neurodegenerative disorders, epilepsy, autism, and ciliopathies (Figure 7E,F, Supplemental Spreadsheet 5). As ciliopathies are often associated with neurodevelopmental defects, we investigated ciliogenesis in the *Ohia* mutant brain and showed a ciliation defect (Figure 7G-I). Overall, these findings point to a role for *Sap130* in the epigenetic regulation of gene expression changes underlying the brain defects in the *Ohia* mutant mice.

To further assess impact of the *Sap130* mutation on DNAm and transcriptional changes in the *Ohia* mutant brain, we examined three-way intersection of the DEGs, the DMR associated genes and genes recovered from the Sap130 ChIPseq analysis. Such three-way intersection of the downregulated DEGs yielded 161 genes (Figure 6B). Pathway enrichment for Mouse Phenotype and Disease terms yielded similar results to that seen in analysis of the DEG and DEG/DMR intersecting genes (Figure 8A,B, Supplemental Spreadsheet 3). This suggests the genes contributing to the neurological diseases and phenotypes are likely targets of Sap130 mediated epigenetic repression. In contrast, three-way intersection of the upregulated DEGs yielded only 43 genes and no significant Mouse Phenotype or Disease terms were recovered (Figure S3). Together these findings suggest Sap130 mediated epigenetic repression of transcription may underlie the transcriptomic changes observed in the *Ohia* mutant mice.

**Figure 8.**
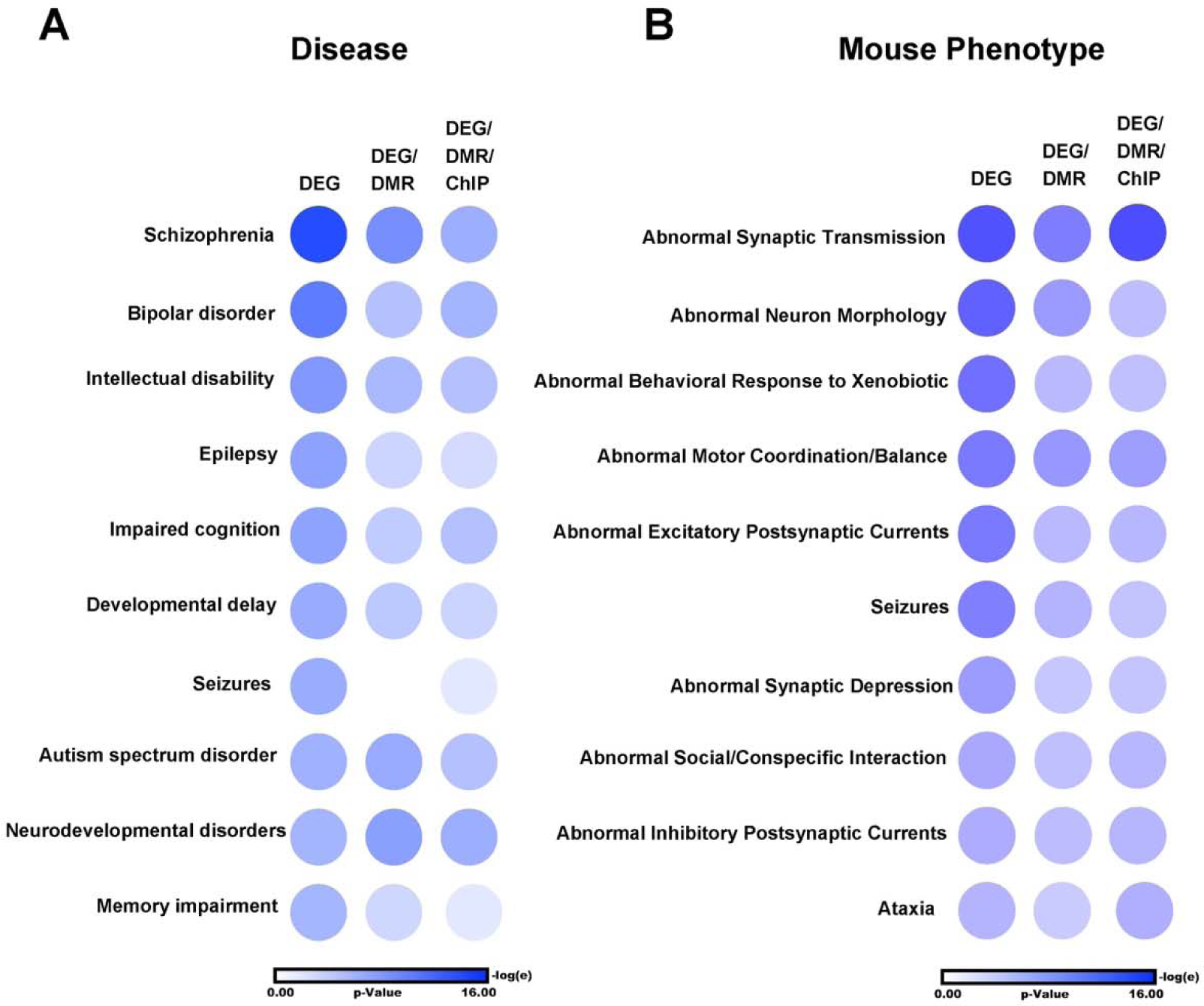
DEGs associated with Disease and Mouse Phenotype are enriched in DMR and Sap130 ChIP binding sites in the *Ohia* mutant brain. **A-B**. Circle plots showing significant Disease (A) and Mouse Phenotype (B) association identified by ToppGene enrichment analysis of the RNASeq recovered down regulated DEGs (first column), genes shared between the down regulated DEGs and DMR associated genes (second column) and the down regulated DEGs, DMR associated genes, and ChIPseq target genes (third column). All three analyses yielded the same Diseases (A) and Mouse Phenotypes (B), but with somewhat different p-value ranking as shown by the color coding in the circle plot.

### Protein-Protein Interactome Network Analysis

To further functionalize the downregulated/differentially methylated genes in the *Ohia* brain, STRING-db was used to construct a protein-protein interaction (PPI) network (Supplementary Spreadsheet 6). The PPI network assembled comprised 132 genes, 82 from the 127 hypomethylated/downregulated genes (FDR=0.05) (Figure S6A) and 50 additional genes recovered by STRING-db (Figure S6A; Supplementary Spreadsheet 6). ToppGene analysis yielded significant terms for Mouse Phenotype, Human Phenotype, Disease, and Pathway. For Human Phenotype, notable was recovery of polymicrogyria (Figure S6B), a defect involving abnormal small cortical folds in the forebrain, a phenotype also reported among human fetuses with HLHS and in primary microcephaly^48,49^. Also relevant to clinical findings in HLHS patients, we observed abnormal hippocampal morphology and multiple seizure related terms (Figure S6B). For Mouse Phenotype, we recovered terms such as cerebral hemisphere morphology, abnormal forebrain morphology, abnormal neuron morphology, and abnormal inhibitory postsynaptic currents (Figure S6C). Disease terms recovered included several also seen with the DMR/DEG analysis, such as neurodevelopmental disorder, autism, impaired cognition, and epilepsy (Figure S6D). Interestingly, Pathway enrichment yielded multiple cilia related terms, with the top pathway being “cargo targeting to cilium”, and also “cargo trafficking to periciliary membrane” and “cilium assembly” (Figure S6E). These findings further point to the disturbance of cilia, supporting the recovery of ciliopathies in the Sap130 ChIPseq analysis, and the ciliation defect observed in the *Ohia* mutant brain.

### Forebrain Specific Deletion of *Sap130* Causes Microcephaly

To investigate cell autonomy of the microcephaly phenotype in the *Ohia^m/m^* mutant mice, we conducted Cre deletion using the forebrain specific Emx1-Cre driver together with a *Sap130* floxed allele. This Cre driver is expressed in excitatory neurons and glia in the developing cerebral cortex from E9.5 onwards^50^. The Emx1-Cre:*Sap130^f/-^*mice are adult viable without CHD and yet they had microcephaly with smaller forebrain (Figure 9A). This was associated with a reduction in the brain to body weight ratio (Figure 9B), suggesting the microcephaly is cell autonomous and unrelated to altered cardiovascular hemodynamics. Unlike the *Ohia* mutant, this phenotype showed complete (100%) penetrance in the Emx1-Cre:*Sap130^f/-^* mice. Real time PCR analysis showed marked reduction in *Sap130* transcripts in the forebrain (Figure 9C). Among the 28 genes known to cause primary microcephaly, many of which are centrosome related, four were up regulated in the Emx1-Cre:*Sap130^f/-^* forebrain (*Wdr62*, *Cit, Ncapd2, Pcnt;* Supplemental Spreadsheet 7), and confirmed by real time PCR analysis (*Wdr62* shown in Figure 9D). We note centrosome amplification can cause spindle defects and is one of the pathogenic mechanisms associated with primary microcephaly^51^. These findings indicate a cell autonomous requirement for *Sap130* in expansion of neural progenitors, deficiency of which likely underlies the microcephaly phenotype observed in *Ohia* mutant mice.

**Figure 9.**
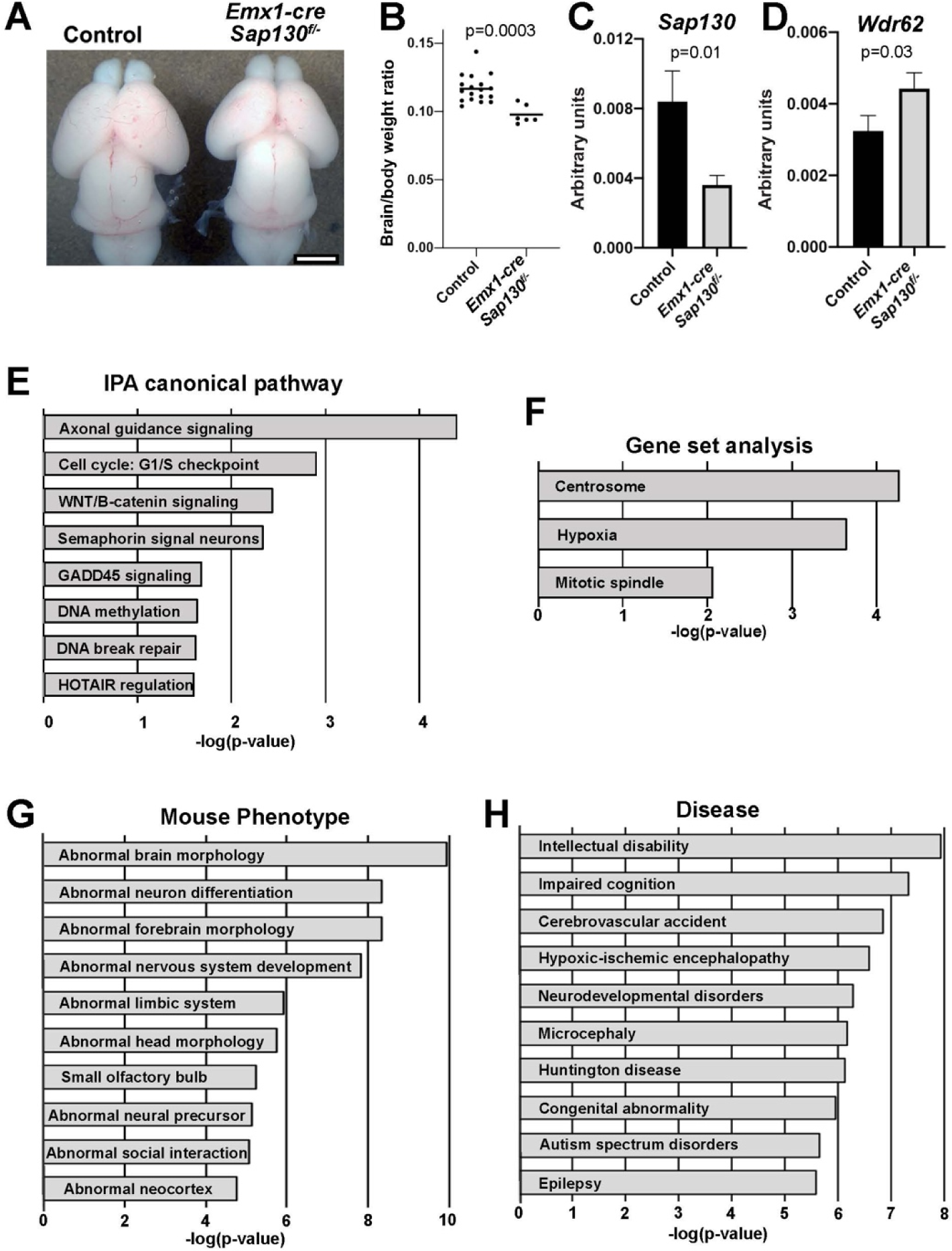
*Emx1-cre Sap130^f/-^* Mice Show Microcephaly With Transcriptome Profiling Showing Abnormal Neurodevelopment. **A.** Wildtype newborn control (left) and *Emx1-cre Sap130^f/-^* mutant (right) brains. Scale bar = 500µm. **B.** Brain/body weight ratio for newborn control and *Emx1-cre Sap130^f/-^* mice confirm brain hypoplasia in the *Emx1-cre Sap130^f/-^* mice. **C.** qPCR of *Sap130* transcripts in E14.5 forebrain tissue from 4 control and 4 *Emx1-cre Sap130^f/-^* mice showed decrease in *Sap130* transcripts, demonstrating efficacy of the Emx1-Cre mediated *Sap130* deletion. **D.** qPCR of *Wdr62* transcripts in E14.5 forebrain tissue from 4 control and 4 *Emx1-cre Sap130^f/-^* mice showed increase transcript expression, indicating possible centrosome amplification. **E-H.** IPA and ToppGene analysis of DEGs from RNAseq of E14.5 forebrain tissue from *Emx1-cre Sap130^f/-^* vs. control mice. Shown are the results obtained with IPA Canonical Pathway (E), Gene Set (F) for hallmark centrosome, hypoxia, and mitotic spindle genes, and ToppGene analysis for Mouse Phenotype (G) and Disease (H).

RNAseq analysis of the E14.5 forebrain of *Emx1-cre*:*Sap130^f/-^* mice (n=3) and littermate controls (n=3) recovered 369 DEGs, 202 down and 167 upregulated (FDR<0.1; Supplemental Spreadsheet 7). IPA pathway enrichment analysis of the DEGs identified axonal guidance as the top pathway, followed by cell cycle, Wnt signaling, and semaphorin signaling in neurons (Figure 9E). Also observed were GADD45 signaling, a stress activated pathway induced by DNA damage and cell cycle checkpoint control, DNA break repair, and HOTAIR regulation - a lncRNA regulating cell cycle and apoptosis via epigenetic gene silencing by providing a scaffold for recruitment of proteins in the Sin3A complex such as REST/Co-REST and lysine demethylase LSD1 (Figure 9E). Gene set enrichment also recovered “Centrosome” and “Mitotic Spindle”, further supporting mitotic spindle defects, and “Hypoxia” was also observed, despite the normal cardiovascular system in the *Emx1-cre*:*Sap130^f/-^* mice (Figure 9F).

Mouse Phenotype terms recovered included abnormal neuronal differentiation, abnormal forebrain morphology, abnormal limbic system, small olfactory bulb, abnormal neural precursor, abnormal social interaction and abnormal neocortex (Figure 9G). For “Disease” enrichment, intellectual disability was recovered as the top pathway, and additionally many Disease terms were recovered that were also seen in the *Ohia* mutant mice such as impaired cognition, neurodevelopmental disorders, microcephaly, and autism (Figure 9H). Cerebrovascular accidents and hypoxic-ischemic encephalopathy were also identified, suggesting a role for hypoxia in the neurological deficits independent of any cardiac defects (Figure 9H). Together these findings suggest microcephaly in *Ohia* mutant mice is not secondary to CHD but may a reflect cell autonomous requirement for *Sap130* in normal forebrain neurodevelopment.

### Adult *Pcdha9^m/m^* and Emx1-Cre Deleted *Sap130* Mice Exhibit Behavioral Deficits

The molecular profiling of the *Ohia^m/m^* fetal brain predicts impaired cognitive function with possible learning/memory defects and autism spectrum disorder. To assess for these neurobehavorial deficits, we generated viable adult mice homozygous for the *Pcdha9* mutation, and mice with forebrain deletion of *Sap130 (*Emx1-Cre:*Sap130^f/-^).* Brain anatomy of the *Emx1-cre:Sap130^f/-^* and *Pcdha9^m/m^* adult mice were analyzed using anatomical brain MRI. The *Emx1-cre:Sap130^f/-^* mice showed significant reduction in total forebrain volume, and volume of the corpus callosum, cortex, and hippocampus (Figure 10A-F). In the *Pcdha9* mutant mice, only mild hippocampal dysplasia was observed but no other appreciable changes in brain structure or size (Figure 10 G,H). Neurobehavioral phenotypes were further assessed using three tests to evaluate spatial learning and memory, fear associative learning, and sociability. These tests were administered on 18 female and 6 male *Pcdha9^m/m^* mice and 6 male and 3 female *Emx1-cre*:*Sap130^f/-^* mice. For the *Pcdha9^m/m^*mice, equal number of sex and age matched C57BL/6J wildtype mice were assessed as controls. For the *Emx1-cre*:*Sap130^f/-^* mice, littermate controls were used comprising Emx1-Cre:*Sap130^f/+^* mice and *Sap130^f/+^* mice without Emx1-cre.

**Figure 10.**
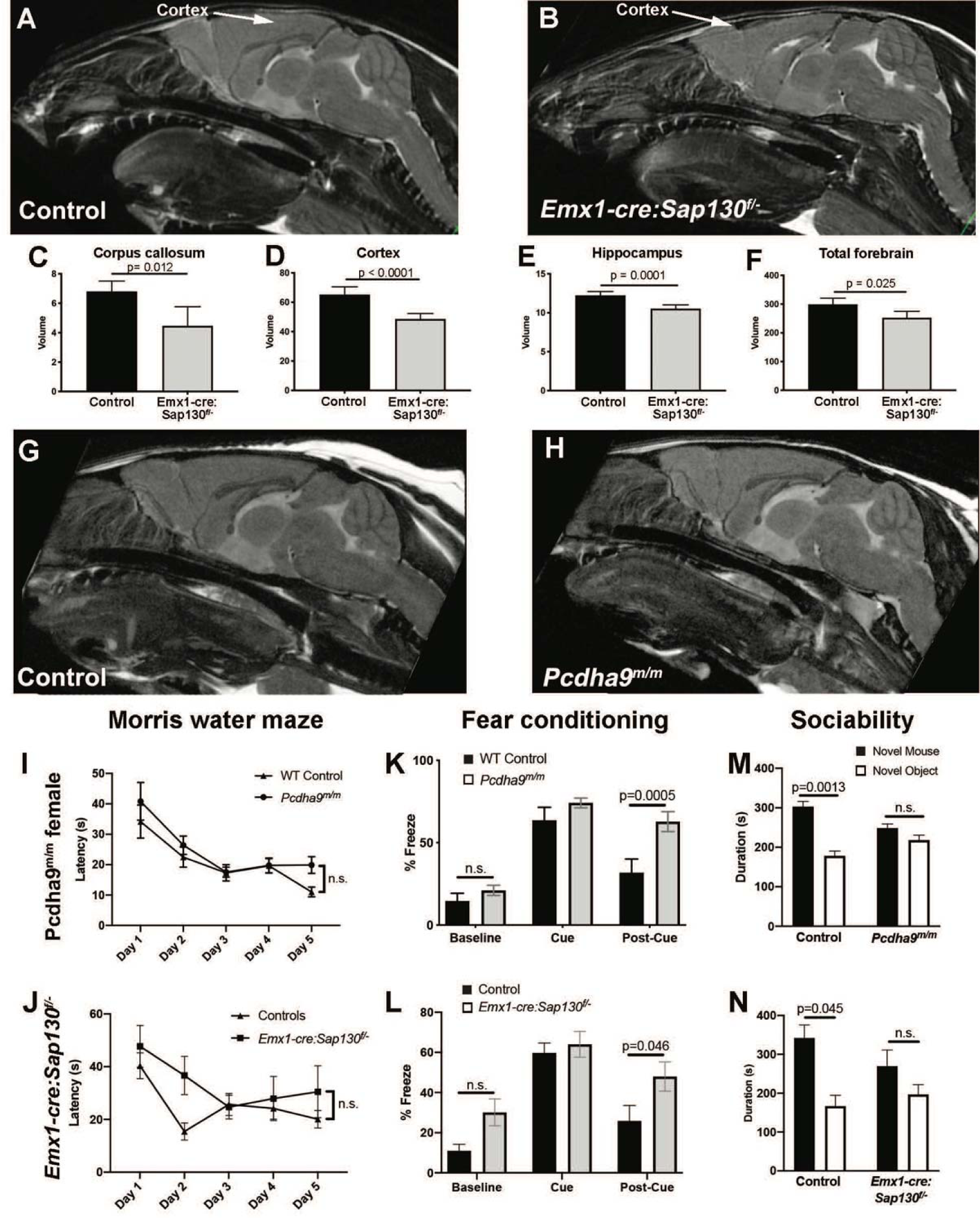
Brain and Behavioral Analysis of *Emx1-cre Sap130^f/-^* and *Pcdha9^m/m^* Adult Mice. **A-F.** Adult brain MRI of wildtype (A), and *Emx1-cre;Sap130^f/-^* adult mice (B). Note the marked reduction in size of the cortex. Quantitative analysis showed significant reduction in brain volumes associated with the corpus callosum (C), the cerebral cortex (D), the hippocampus (E), and forebrain (F). **G,H.** Adult brain MRI of wildtype (A), and *Pcdha9^m/m^*adult mice (B). No change was observed in anatomical structure of the *Pcdha9^m/m^* mouse brain. **I-N.** Neurobehavioral testing was conducted using three tests: Morris water maze (I,J), cued fear conditioning (K,L), and three chamber sociability test (M,N). In both *Pcdha9^m/m^*female mice and *Emx1-cre:Sap130^f/-^* mice, significant deficits were observed for fear conditioning and sociability, but no difference was observed in the Morris water maze. Similar analysis of *Pcdha9^m/m^* male mice showed no change relative to wildtype control male mice (Supplemental Figure S7). Morris water maze analyzed by 2-way repeated measures ANOVA, fear conditioning analyzed by 2-way repeated measures ANOVA, and sociability tested analyzed by two-way and three-way ANOVA.

Spatial learning and memory were interrogated using the Morris water maze whereby mice were trained to find a hidden platform in a pool of water. After training, no significant difference in the time (latency) required to find the hidden platform was observed for either the *Pcdha9^m/m^*or Emx1-Cre:*Sap130*^f/-^ mice vs. controls (Figure 10 I,J, Figures S7,S8). In the fear conditioning test, mice learn to associate a cue (a tone) with an adverse event (foot shock) administered after the cue. After training, mice typically freeze upon hearing the cue, but return to normal activity post cue. Emx1-Cre:*Sap130*^f/-^ mice and *Pcdha9^m/m^* female, but not male mice, showed increased post-cue freezing, indicating impaired associative learning (Figure 10K,L; Figures S7,S8). We also conducted sociability testing in which mice are presented with a novel object vs. a novel mouse. While wildtype and littermate control mice showed preference for the novel mouse (Figure 10M,N), Emx1-Cre:*Sap130^f/-^*mice and *Pcdha9^m/m^* female mice, but not male mice, showed equal time spent exploring the novel object and novel mouse (Figure 10M,N, Figures S7,S8), indicating social interaction deficits. It was not possible to assess gender effects in the Emx1-Cre*:Sap130^f/-^*mice given the smaller cohort size.

## DISCUSSION

CHD patients have increased risk for adverse neurodevelopmental outcome that can significantly degrade their quality of life. While clinical studies have shown this is associated with brain abnormalities and neurobehavioral impairment, the underlying mechanisms remain unclear. Here our studies of the *Ohia* mouse model of HLHS show the same two mutations causing HLHS also can cause brain and neurodevelopmental abnormalities. This is congruent with the fact that many genes highly expressed in the heart are also highly expressed in the brain^12^. Moreover, de novo mutations linked to CHD are enriched for genes known to cause neurodevelopmental defects^12^.

We showed *Ohia* fetal mice can exhibit brain defects comprising forebrain hypoplasia, with severe phenotype comprising overt microcephaly and milder phenotype with modest to no obvious forebrain hypoplasia. Consistent with our findings indicating a developmental etiology, brain volume reduction has been reported with in utero imaging studies of human fetuses with HLHS^52-54^. As the volume reduction is correlated with lower brain oxygenation, this suggests possible disruption of neurovascular coupling - a brain sparing cerebral autoregulatory mechanism that shunts blood to the brain during hypoxia^41,48^. This is supported by clinical studies with in utero ultrasound imaging of fetuses with HLHS^,49^. The recovery of hypoxemic encephalopathy in transcriptome profiling of the Emx1-Cre:*Sap130^f/-^* mouse brain without CHD supports the disturbance of neurovascular coupling independent of altered cardiovascular hemodynamics. Previous studies with blood oxygen level dependent (BOLD) brain MRI in CHD patients in fact showed impaired neurovascular coupling^55^. This was associated with low NO, which regulates vasodilation and blood flow, and also impaired cognitive function^55^.

A cortical neurogenesis defect was observed with deficiency of later born neurons in cortical layers II-IV. This is likely due to the loss of intermediate neural progenitors in the subventricular zone. A cortical neurogenesis defect also has been reported clinically in HLHS fetuses, with volume reduction noted in the cortical subplate and intermediate and ventricular zones^56^. In the *Ohia* mutant brain, the neurogenesis defect is accompanied by the disturbance of REST, a master transcriptional regulator that recruits HDACs to the Sin3A complex to epigenetically regulate neural progenitor expansion and neural stem cell maintenance^33,34,57,58^. Microcephaly with loss of intermediate progenitors has been observed with disturbance of REST via HDAC deficiency and also with mutations in *ZNF335,* another component of the Sin3A complex required for REST regulation of neurogenesis^59,60^.

We further showed the loss of intermediate progenitors likely arises from a centrosome defect causing multipolar spindle formation and metaphase block with ensuing apoptosis. Mutations in centrosome genes and spindle defects are commonly observed in primary microcephaly, with *Wdr62* being one of the major genetic causes of primary microcephaly^61^. *Wdr62* mutations have been implicated in autism, and neuronal deletion of *Wdr62* in mice caused impaired learning/memory and social interaction deficits without microcephaly, suggesting additional roles for the centrosome in neurodevelopment^62^. Interestingly, CHD patients with rare variants in *Wdr62* have been found to have multipolar spindles associated with a cardiomyocyte proliferation defect, indicating the same gene may cause both brain and heart defects^63^. The expression of *Wdr62* and several other microcephaly associated centrosome genes were elevated in *Ohia* mutant brain, suggesting centrosome amplification, a mechanism also implicated in primary microcephaly^51,64^. Our gene set enrichment analysis recovered “centrosome” and “mitotic spindle” from the Emx1-Cre:*Sap130^f/-^* transcriptome analysis, and “cell cycle” and “centrosome” from the Sap130 ChIPseq analysis. Together these findings suggest *Sap130* may regulate centrosome function critical for cortical neurogenesis.

Interestingly, Sap130 ChIPseq analysis recovered “ciliopathy”. We observed a ciliation defect in the *Ohia* mutant brain. Multiple cilia-related pathways were also recovered in our PPI network analysis. As cilia assembly is templated on a basal body derived from the centrosome, centrosome defects can contribute to both ciliation and mitotic spindle defects. We note HLHS patients have been reported to have increased burden for ciliopathy related variants, and conversely patients with various ciliopathies have been reported to have HLHS^65^. As both HLHS and ciliopathies are rare birth defects, their cooccurrence would suggest a mechanistic link. This could involve the role of cilia in neural circuit formation and the regulation of neuronal connectivity^66^. The finding of holoprosencephaly among *Ohia* mutants and also HLHS patients would also suggest possible disturbance of cilia transduced hedgehog signaling^67^.

The transcriptome profiling indicated the disturbance of CREB signaling, a cAMP and calcium responsive transcriptional pathway regulating neuronal plasticity, long term potentiation and memory encoding. Thus, CREB disturbance could lead to cognitive impairment and memory related diseases such as Alzhemier’s^38^. We note HLHS and other single ventricle CHD patients are reported to have hazards ratio of 2.6 (95% CI:1.8-3.8) for early onset dementia^38,68^. CREB signaling also regulates dopaminergic neurons implicated in autism, with CREB disturbance seen in Timothy syndrome associated with high penetrance for autism^69^. Our transcriptome profiling recovered autism, intellectual disability, impaired cognition, all neurobehavioral deficits similar to those seen in HLHS. This is supported by the results of neurobehavioral assessments showing autism-like social interaction deficits and impaired learning/memory in both the Emx1-Cre:*Sap130^f/-^* mice with microcephaly, and the *Pcdha9* mutant mice with normal brain anatomy. This suggests *Sap130* and the *Pcdha* gene cluster may act on convergent processes or pathways. While genes from the *Pcdha* gene cluster provide unique cell surface barcodes defining neuronal identify that regulate neural circuit assembly, expression of the *Pcdha* gene cluster involves epigenetic regulation mediated by the Sin3A complex^70,71^. This may underlie the similar neurobehavioral findings of the Emx1-Cre:*Sap130^f/-^* and *Pcdha9* mutant mice.

The neurological phenotypes and diseases recovered from the transcriptome profiling in the *Ohia* and Emx1-Cre:*Sap130^f/-^*mutant mice were associated with downregulated DEGs. As these down regulated DEGs were also differentially methylated and are targets of Sap130 binding, this suggests Sap130 may epigenetically regulate gene expression. We note transcription from the *Pcdha* gene cluster and other genes with the CTCF motif requires promoter demethylation with recruitment of TET demethylases via the Sin3A complex^72,73^. Thus, we propose promoter demethylation by the Sin3A complex may be suppressed with Sap130 deficiency, thus contributing to the observed downregulation of gene expression in *Ohia* mutant mice. As the *Ohia* Sap130 allele is hypomorphic, this may explain the incomplete penetrance for microcephaly in the *Ohia* mutant mice, while complete penetrance for microcephaly in the *Emx1-Cre:Sap130^f/-^*mice may be explained by the loss of function orchestrated by the Cre deletion. Such epigenetic perturbation may account for the finding of similar neurobehavioral phenotypes in the *Pcdha9^m/m^ and Emx1-Cre:Sap130^f/-^*mice. We note disturbance of DNAm in the clustered protocadherins has been associated with various neurodevelopmental disorders, including William syndrome with aortic valve defects^74^. Patients with William syndrome exhibit DNA hypermethylation and are described to have hypermethylation of *Pcdha9* and other genes in the clustered protocadherins^75^. The recent identification of DNAm signatures for over 40 neurodevelopmental disorders, including Witteveen-Kolk syndrome with Sin3A mutations, further support the involvement of DNAm disturbance in neurodevelopmental disorders^76^.

Overall, our study uncovered cell autonomous and nonautonomous defects together with epigenetic dysregulation contributing to the neurodevelopmental defects in the *Ohia* HLHS mouse model. The mechanistic insights obtained support new avenues for therapy to improve neurodevelopmental outcome, such as with manipulation of the epigenome via diet or pharmacological intervention^77^. It is worth noting as mice with microcephaly or normal brain anatomy can have similar neurobehavioral deficits, more sophisticated functional brain imaging may be needed to fully assess the risk for adverse neurodevelopmental outcome. We expect these findings will have broad clinical relevance beyond HLHS, as adverse neurodevelopmental outcome is observed among CHD of a broad spectrum.

## METHODS

Extended, more detailed methods are provided in the Supplemental Material. All data supporting the findings of this study are available from the corresponding author upon reasonable request.

### Mouse Husbandry

Mouse studies were conducted under a University of Pittsburgh Institutional Animal Care and Use Committee approved animal study protocol. *Ohia* and *Pcdha9* (c.2389_2399del; [p.Asp796Phefs∗]) mice generated by CRISPR gene editing were maintained in the C57BL/6J background. B6.129S2-Emx1tm1(cre)Krj/J mice with *Emx1-cre* driver were purchased from Jackson Laboratory (Strain #005628) and intercrossed with *Sap130^f/f^*and *Sap130^+/-^* mice generated previously^20^.

### Immunostaining and Confocal Microscopy

*Ohia^m/m^* fetuses were fixed in 4% PFA overnight, and cryoembedded. Cryosections were generated and stained with different antibodies to various neural markers (see Supplemental Methods).

### Mouse Embryonic Fibroblast Analysis

Mouse embryonic fibroblasts (MEFs) were isolated from E14.5-E15.5 embryos as previously described, with three independent MEF lines analyzed. Cells were plated on glass coverslips, fixed in 4% PFA, stained with antibodies to α-tubulin (Abcam; ab15246), and γ-tubulin (Sigma; T6557) and counterstained with DAPI (Thermo Fisher Scientific; D1306).

### Histological Reconstructions Using Episcopic Confocal Microscopy

E14.5 or newborn mice were euthanized, the heads embedded in paraffin for episcopic confocal microscopy (ECM) as previously described^20^. The 2D serial ECM image stacks collected are digitally resliced and also 3D reconstructed using the OsiriX Dicom viewer (https://www.osirix-viewer.com) to assess brain anatomical structures.

### RNA Sequencing

RNA sequencing of brain tissue was conducted using standard protocols on Illumina HiSeq 2000 (BGI Americas) with 100-bp paired-end reads, and using standard bioinformatics pipeline (see Supplemental Methods) and aligned to mm10 (NCBI build 38). Differential expression analyses were performed with edgeR, and differentially expressed genes for *Ohia^m/m^* mutant brain were identified with FDR ≤0.05 (Benjamini–Hochberg), and for *Emx1-cre*:*Sap130^f/-^* with FDR<0.1. Pathway analysis was carried out genes identified to be differentially expressed.

### Chromatin Immunoprecipitation Sequencing

Sap130 chromatin immunoprecipitation was performed with rabbit anti-Sap130 antibody (A302-491A, Bethyl laboratories) and an iDeal ChIP–seq Kit for Transcription Factors (Diagenode) as previously described^20^ (see Supplemental Methods). Reads were aligned to mm10 and Sap130 target regions were identified with MACS1.4.2. Sap130 binding sites situated within 1 kb of transcriptional start sites were used to recover genes likely to be targets of Sap130 epigenetic regulation. Motif enrichment analysis was performed using MEME suite version 5.4.1.

### Methylation Analysis

Methylation analysis was conducted using the Illumina Infinium mouse methylation beadchip using standard protocol (see Supplemental Methods). Probes with FDR≤0.1 were considered significantly differentially methylated. Differentially methylated regions (DMRs) were identified using the DMRcate (v 2.4.1) with FDR of 0.05.

### Mouse Behavioral Phenotyping

Mouse behavior was assessed using three training paradigms: the Morris water maze, fear conditioning, and sociability. Testing was completed by operator blinded to genotype. See Supplemental Methods for more details.

### Magnetic Resonance Imaging

Following behavioral testing, mice underwent in vivo brain MRI carried out using a Bruker BioSpec 70/30 USR spectrometer (Bruker BioSpin MRI, Billerica, MA, USA) operating at 7-Tesla with quadrature radio-frequency volume coil with inner-diameter of 35 mm.

### Statistical Analysis

Data were analyzed using GraphPad Prism 9 (GraphPad Software). For cell quantification independent samples T-test was used (*p<0.05). For mouse behavioral analysis, data were analyzed by two-way, two-way repeated measures, or three-way analysis of variance (ANOVA).

## Author Contributions

GCG, BJG, MGS performed mouse husbandry and genotyping, GCG performed mouse brain immunostaining and cell culture work; GCG, AB, LH, HY performed RNAseq, ChIPseq, and methylation experiments and analysis; MG assisted with PPI visualization, GCG and DS performed mouse behavioral testing and analysis; GCG, WTR performed episcopic confocal microscopy analysis of mouse brains; MCS and YLW performed brain MRI imaging and analysis, GCG and CWL participated in the experimental design and analysis of results; GCG, BJG, AP, CWL performed manuscript writing and editing. All authors approved the final manuscript.

## REFERENCES

1. Hoffman, J.I. & Kaplan, S. The incidence of congenital heart disease. Journal of the American college of cardiology 39, 1890–1900 (2002).

2. Marino, B.S. et al. Neurodevelopmental outcomes in children with congenital heart disease: evaluation and management: a scientific statement from the American Heart Association. Circulation 126, 1143–1172 (2012).

3. Marelli, A., Miller, S.P., Marino, B.S., Jefferson, A.L. & Newburger, J.W. Brain in Congenital Heart Disease Across the Lifespan. Circulation 133, 1951–1962 (2016).

4. Gaynor, J.W. et al. Patient characteristics are important determinants of neurodevelopmental outcome at one year of age after neonatal and infant cardiac surgery. The Journal of thoracic and cardiovascular surgery 133, 1344–1353. e3 (2007).

5. Fuller, S. et al. Predictors of impaired neurodevelopmental outcomes at one year of age after infant cardiac surgery. European journal of cardio-thoracic surgery 36, 40–48 (2009).

6. Newburger, J.W. et al. Early developmental outcome in children with hypoplastic left heart syndrome and related anomalies: the single ventricle reconstruction trial. Circulation 125, 2081–2091 (2012).

7. Hinton, R.B. et al. Prenatal head growth and white matter injury in hypoplastic left heart syndrome. Pediatric research 64, 364–369 (2008).

8. Miller, S.P. et al. Abnormal brain development in newborns with congenital heart disease. New England Journal of Medicine 357, 1928–1938 (2007).

9. Owen, M., Shevell, M., Majnemer, A. & Limperopoulos, C. Abnormal brain structure and function in newborns with complex congenital heart defects before open heart surgery: a review of the evidence. Journal of Child Neurology 26, 743–755 (2011).

10. Glauser, T.A., Rorke, L.B., Weinberg, P.M. & Clancy, R.R. Congenital brain anomalies associated with the hypoplastic left heart syndrome. Pediatrics 85, 984–990 (1990).

11. Zaidi, S. et al. De novo mutations in histone-modifying genes in congenital heart disease. Nature 498, 220–223 (2013).

12. Homsy, J. et al. De novo mutations in congenital heart disease with neurodevelopmental and other congenital anomalies. Science 350, 1262–1266 (2015).

13. Schanen, N.C. Epigenetics of autism spectrum disorders. Human molecular genetics 15, R138–R150 (2006).

14. Bucholz, E.M. et al. Socioeconomic Status and Long-term Outcomes in Single Ventricle Heart Disease. Pediatrics 146(2020).

15. Goldberg, C.S. et al. Behavior and Quality of Life at 6 Years for Children With Hypoplastic Left Heart Syndrome. Pediatrics 144(2019).

16. Sananes, R. et al. Six-Year Neurodevelopmental Outcomes for Children With Single-Ventricle Physiology. Pediatrics 147(2021).

17. Siciliano, R.E. et al. Cognitive Function in Pediatric Hypoplastic Left Heart Syndrome: Systematic Review and Meta-Analysis. J Pediatr Psychol 44, 937–947 (2019).

18. Hangge, P.T. et al. Microcephaly is associated with early adverse neurologic outcomes in hypoplastic left heart syndrome. Pediatr Res 74, 61–7 (2013).

19. Laraja, K. et al. Neurodevelopmental Outcome in Children after Fetal Cardiac Intervention for Aortic Stenosis with Evolving Hypoplastic Left Heart Syndrome. J Pediatr 184, 130–136 e4 (2017).

20. Liu, X. et al. The complex genetics of hypoplastic left heart syndrome. Nature genetics 49, 1152–1159 (2017).

21. Witteveen, J.S. et al. Haploinsufficiency of MeCP2-interacting transcriptional co-repressor SIN3A causes mild intellectual disability by affecting the development of cortical integrity. Nature Genetics 48, 877–887 (2016).

22. Amir, R.E. et al. Rett syndrome is caused by mutations in X-linked MECP2, encoding methyl-CpG-binding protein 2. Nat Genet 23, 185–8 (1999).

23. Shirvani-Farsani, Z., Maloum, Z., Bagheri-Hosseinabadi, Z., Vilor-Tejedor, N. & Sadeghi, I. DNA methylation signature as a biomarker of major neuropsychiatric disorders. Journal of Psychiatric Research 141, 34–49 (2021).

24. Aref-Eshghi, E. et al. Evaluation of DNA Methylation Episignatures for Diagnosis and Phenotype Correlations in 42 Mendelian Neurodevelopmental Disorders. Am J Hum Genet 106, 356–370 (2020).

25. Zhu, F. et al. Sin3a-Tet1 interaction activates gene transcription and is required for embryonic stem cell pluripotency. Nucleic Acids Res 46, 6026–6040 (2018).

26. Thu, C.A. et al. Single-cell identity generated by combinatorial homophilic interactions between α, β, and γ protocadherins. Cell 158, 1045–1059 (2014).

27. Mountoufaris, G. et al. Multicluster Pcdh diversity is required for mouse olfactory neural circuit assembly. Science 356, 411–414 (2017).

28. Teekakirikul, P., et al. Common deletion variants causing protocadherin-alpha deficiency contribute to the complex genetics of BAV and left-sided congenital heart disease. HGG Adv 2 (2021).

29. Uctepe, E. et al. A Witteveen-Kolk Syndrome Patient with Reflux Disease and a de novo Deletion of the SIN3A Gene. Molecular Syndromology, 1–6 (2024).

30. Anitha, A. et al. Protocadherin α (PCDHA) as a novel susceptibility gene for autism. Journal of Psychiatry and Neuroscience 38, 192–198 (2013).

31. Nicholas, A.K. et al. The molecular landscape of ASPM mutations in primary microcephaly. Journal of medical genetics 46, 249–253 (2009).

32. Nicholas, A.K. et al. WDR62 is associated with the spindle pole and is mutated in human microcephaly. Nature genetics 42, 1010–1014 (2010).

33. Gao, Z. et al. The master negative regulator REST/NRSF controls adult neurogenesis by restraining the neurogenic program in quiescent stem cells. Journal of Neuroscience 31, 9772–9786 (2011).

34. Ballas, N., Grunseich, C., Lu, D.D., Speh, J.C. & Mandel, G. REST and its corepressors mediate plasticity of neuronal gene chromatin throughout neurogenesis. Cell 121, 645–657 (2005).

35. Rodenas-Ruano, A., Chávez, A.E., Cossio, M.J., Castillo, P.E. & Zukin, R.S. REST-dependent epigenetic remodeling promotes the developmental switch in synaptic NMDA receptors. Nature neuroscience 15, 1382–1390 (2012).

36. Bray, N.J. et al. Haplotypes at the dystrobrevin binding protein 1 (DTNBP1) gene locus mediate risk for schizophrenia through reduced DTNBP1 expression. Human molecular genetics 14, 1947–1954 (2005).

37. Lang, F., Strutz-Seebohm, N., Seebohm, G. & Lang, U.E. Significance of SGK1 in the regulation of neuronal function. The Journal of physiology 588, 3349–3354 (2010).

38. Saura, C.A. & Valero, J. The role of CREB signaling in Alzheimer’s disease and other cognitive disorders. Rev Neurosci 22, 153–69 (2011).

39. Gau, D. et al. Phosphorylation of CREB Ser142 regulates light-induced phase shifts of the circadian clock. Neuron 34, 245–53 (2002).

40. Stackhouse, T.L. & Mishra, A. Neurovascular Coupling in Development and Disease: Focus on Astrocytes. Front Cell Dev Biol 9, 702832 (2021).

41. Donofrio, M. et al. Autoregulation of cerebral blood flow in fetuses with congenital heart disease: the brain sparing effect. Pediatric cardiology 24, 436–443 (2003).

42. Riccio, A. et al. A nitric oxide signaling pathway controls CREB-mediated gene expression in neurons. Mol Cell 21, 283–94 (2006).

43. Williams, K. et al. TET1 and hydroxymethylcytosine in transcription and DNA methylation fidelity. Nature 473, 343–348 (2011).

44. Ávila-Mendoza, J., Subramani, A. & Denver, R.J. Krüppel-like factors 9 and 13 block axon growth by transcriptional repression of key components of the cAMP signaling pathway. Frontiers in Molecular Neuroscience, 217 (2020).

45. Jiang, J. et al. A core Klf circuitry regulates self-renewal of embryonic stem cells. Nature cell biology 10, 353–360 (2008).

46. Lavallée, G. et al. The Kruppel-like transcription factor KLF13 is a novel regulator of heart development. The EMBO journal 25, 5201–5213 (2006).

47. Dekker, R.J. et al. Endothelial KLF2 links local arterial shear stress levels to the expression of vascular tone-regulating genes. Am J Pathol 167, 609–18 (2005).

48. Kinnear, C. et al. Abnormal fetal cerebral and vascular development in hypoplastic left heart syndrome. Prenat Diagn 39, 38–44 (2019).

49. Bhat, V. et al. Mutations in WDR62, encoding a centrosomal and nuclear protein, in Indian primary microcephaly families with cortical malformations. Clin Genet 80, 532–40 (2011).

50. Gorski, J.A. et al. Cortical excitatory neurons and glia, but not GABAergic neurons, are produced in the Emx1-expressing lineage. Journal of Neuroscience 22, 6309–6314 (2002).

51. Zhang, Y. et al. Overexpression of WDR62 is associated with centrosome amplification in human ovarian cancer. Journal of ovarian research 6, 1–6 (2013).

52. Sadhwani, A. et al. Fetal Brain Volume Predicts Neurodevelopment in Congenital Heart Disease. Circulation 145, 1108–1119 (2022).

53. Limperopoulos, C. et al. Brain volume and metabolism in fetuses with congenital heart disease: evaluation with quantitative magnetic resonance imaging and spectroscopy. Circulation 121, 26–33 (2010).

54. Sun, L. et al. Reduced fetal cerebral oxygen consumption is associated with smaller brain size in fetuses with congenital heart disease. Circulation 131, 1313–23 (2015).

55. Schmithorst, V.J. et al. Impaired Neurovascular Function Underlies Poor Neurocognitive Outcomes and Is Associated with Nitric Oxide Bioavailability in Congenital Heart Disease. Metabolites 12(2022).

56. Rollins, C.K. et al. Regional Brain Growth Trajectories in Fetuses with Congenital Heart Disease. Ann Neurol 89, 143–157 (2021).

57. Qureshi, I.A., Gokhan, S. & Mehler, M.F. REST and CoREST are transcriptional and epigenetic regulators of seminal neural fate decisions. Cell Cycle 9, 4477–86 (2010).

58. Huang, Y., Myers, S.J. & Dingledine, R. Transcriptional repression by REST: recruitment of Sin3A and histone deacetylase to neuronal genes. Nature neuroscience 2, 867–872 (1999).

59. Tang, T. et al. HDAC1 and HDAC2 regulate intermediate progenitor positioning to safeguard neocortical development. Neuron 101, 1117–1133. e5 (2019).

60. Yang, Y.J. et al. Microcephaly gene links trithorax and REST/NRSF to control neural stem cell proliferation and differentiation. Cell 151, 1097–1112 (2012).

61. Chen, J.-F. et al. Microcephaly disease gene Wdr62 regulates mitotic progression of embryonic neural stem cells and brain size. Nature communications 5, 1–13 (2014).

62. Xu, D. et al. WDR62-deficiency Causes Autism-like Behaviors Independent of Microcephaly in Mice. Neurosci Bull 39, 1333–1347 (2023).

63. Hao, L. et al. WDR62 variants contribute to congenital heart disease by inhibiting cardiomyocyte proliferation. Clin Transl Med 12, e941 (2022).

64. Marthiens, V. et al. Centrosome amplification causes microcephaly. Nature cell biology 15, 731–740 (2013).

65. Geddes, G.C., Stamm, K., Mitchell, M., Mussatto, K.A. & Tomita-Mitchell, A. Ciliopathy variant burden and developmental delay in children with hypoplastic left heart syndrome. Genetics in Medicine 19, 711–714 (2017).

66. Guo, J. et al. Primary Cilia Signaling Shapes the Development of Interneuronal Connectivity. Dev Cell 42, 286–300 e4 (2017).

67. Nanni, L. et al. The Mutational Spectrum of the Sonic Hedgehog Gene in Holoprosencephaly: SHH Mutations Cause a Significant Proportion of Autosomal Dominant Holoprosencephaly. Human Molecular Genetics 8, 2479–2488 (1999).

68. Bagge, C.N. et al. Risk of Dementia in Adults With Congenital Heart Disease: Population-Based Cohort Study. Circulation 137, 1912–1920 (2018).

69. Servili, E., Trus, M., Sajman, J., Sherman, E. & Atlas, D. Elevated basal transcription can underlie timothy channel association with autism related disorders. Prog Neurobiol 191, 101820 (2020).

70. Canzio, D. et al. Antisense lncRNA transcription mediates DNA demethylation to drive stochastic protocadherin α promoter choice. Cell 177, 639–653. e15 (2019).

71. Canzio, D. & Maniatis, T. The generation of a protocadherin cell-surface recognition code for neural circuit assembly. Curr Opin Neurobiol 59, 213–220 (2019).

72. Salas-Armenteros, I. et al. Human THO-Sin3A interaction reveals new mechanisms to prevent R-loops that cause genome instability. EMBO J 36, 3532–3547 (2017).

73. Grunseich, C. et al. Senataxin Mutation Reveals How R-Loops Promote Transcription by Blocking DNA Methylation at Gene Promoters. Mol Cell 69, 426–437 e7 (2018).

74. El Hajj, N., Dittrich, M. & Haaf, T. Epigenetic dysregulation of protocadherins in human disease. Semin Cell Dev Biol 69, 172–182 (2017).

75. Strong, E. et al. Symmetrical dose-dependent DNA-methylation profiles in children with deletion or duplication of 7q11. 23. The American Journal of Human Genetics 97, 216-227 (2015).

76. Coenen-van der Spek, J., et al. DNA methylation episignature for Witteveen-Kolk syndrome due to SIN3A haploinsufficiency. Genet Med 25, 63–75 (2023).

77. Rasmi, Y. et al. The role of DNA methylation in progression of neurological disorders and neurodegenerative diseases as well as the prospect of using DNA methylation inhibitors as therapeutic agents for such disorders. IBRO Neurosci Rep 14, 28–37 (2023).

