## Supplemental Information for "Mitotic Block and Epigenetic Repression Underlie Neurodevelopmental Defects and Neurobehavioral Deficits in Congenital Heart Disease"

**SUPPLEMENTARY INFORMATION**

**Contents:**

**Supplemental Figure S1.** Craniofacial defects in embryonic *Ohia* mice.

**Supplemental Figure S2.** IPA pathway analysis of differentially expressed genes in the *Ohia* mutant brain showed perturbation of neurovascular coupling.

**Supplemental Figure S3.** Integrated Analysis of Genes Shared in Common Among the RNA-seq Upregulated Genes, Genes Associated with DMRs, and Sap130 ChIP-seq Target Genes.

**Supplemental Figure S4.** DNA methylation analysis of *Ohia* mouse brain in mutant mice with severe vs. mild brain phenotype.

**Supplemental Figure S5.** Comparison of disease pathways recovered from analysis of differentially methylated regions in brain tissue of *Ohia* mutant mice with severe vs. mild brain defects.

**Supplemental Figure S6.** Protein-Protein Interactome Network Analysis

**Supplemental Figure S7.** Behavioral Testing of *Pcdha9^m/m^* and EMX1-cre:*Sap130^f/-^* female mice.

**Supplemental Figure S8.** Behavioral Testing of *Pcdha9^m/m^* Male Mice.

**Supplemental Methods.**

**SUPPLEMENTAL FIGURES**

**Figure S1. Craniofacial defects in embryonic *Ohia* mice.**

*Ohia* mutant mice double homozygous for the Sap130/Pcdha9 mutations exhibit craniofacial abnormalities of varying severity. Shown is a wildtype control E14.5 mouse, and three double homozygous E14.5 *Ohia* mutant mice. The mutant mice exhibit varying degrees of micrognathia including shortened snout with recessed lower jaw (b,c), agnathia with missing lower jaw (d), small head (c,d), low set ears and eye defects (b,c,d).

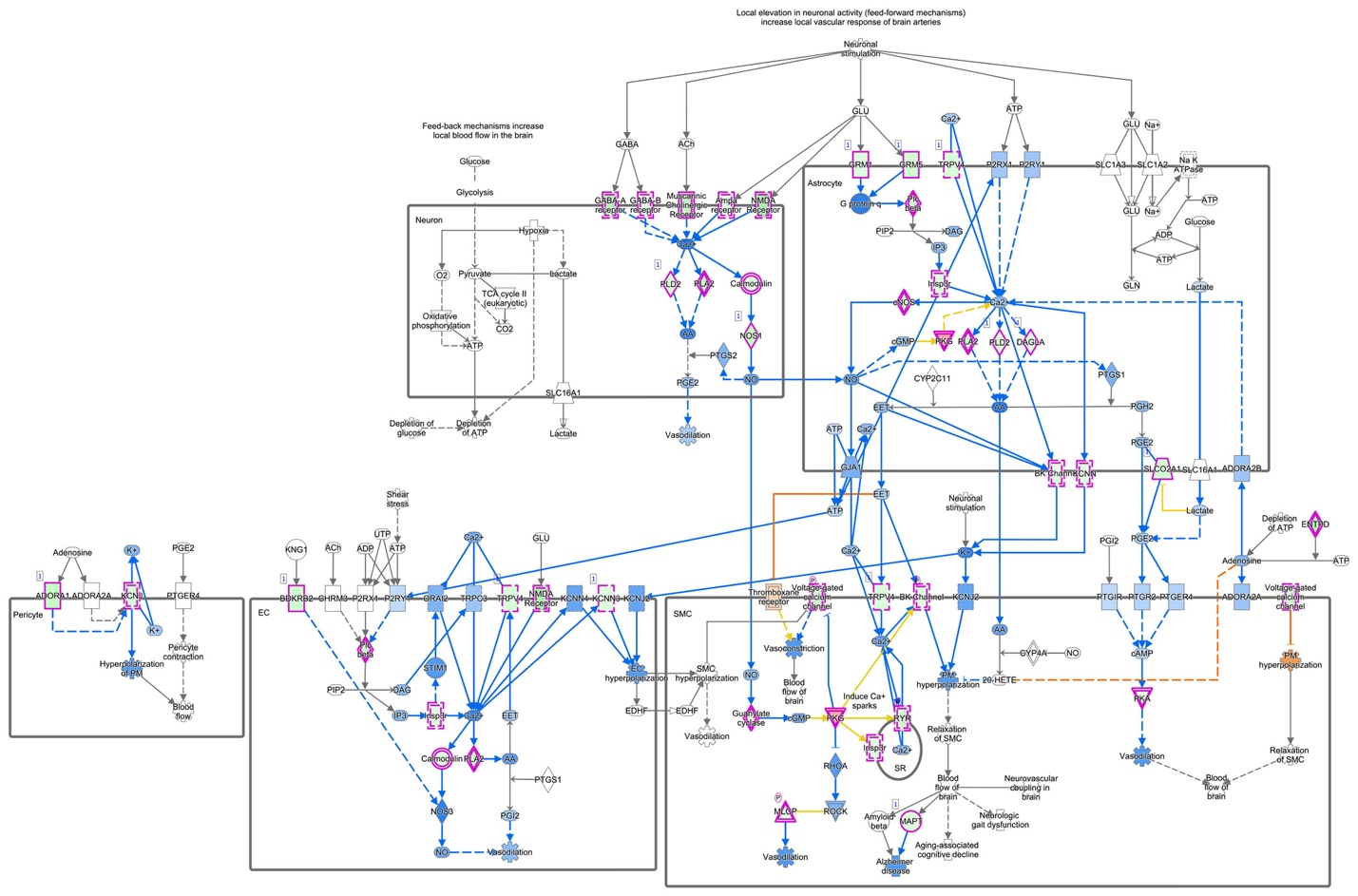

**Figure S2. IPA pathway analysis of differentially expressed genes in the *Ohia* mutant brain showed perturbation of neurovascular coupling.**

IPA analysis of DEGs recovered from the *Ohia* mutant brain recovered neurovascular coupling, a pathway depicted in the IPA diagram above. The purple outlined geometric shapes denote genes differentially expressed, with green fill indicating down regulated expression, and red fill indicating up regulated expression. The individual rectangles correspond to different cell types in the brain including neurons, astrocytes, endothelial cells, smooth muscle cells, and pericytes, and their interactions.

**Figure S3. Integrated Analysis of Genes Shared in Common Among the RNA-seq Upregulated Genes, Genes Associated with DMRs, and Sap130 ChIP-seq Target Genes.**

A. Intersection of RNA-seq upregulated genes and genes identified in the DMRs.
B. Three-way intersection of RNA-seq upregulated genes, genes associated with the DMRs, and Sap130 target genes identified in the ChIP-seq analysis.
C. Toppgene Pathway analysis of 61 genes identified to be transcriptionally up-regulated and also differentially methylated in the *Ohia* brain.
D. Toppgene Pathway analysis of 43 genes identified to be up-regulated, differentially methylated, and also Sap130 target genes in the *Ohia* brain.

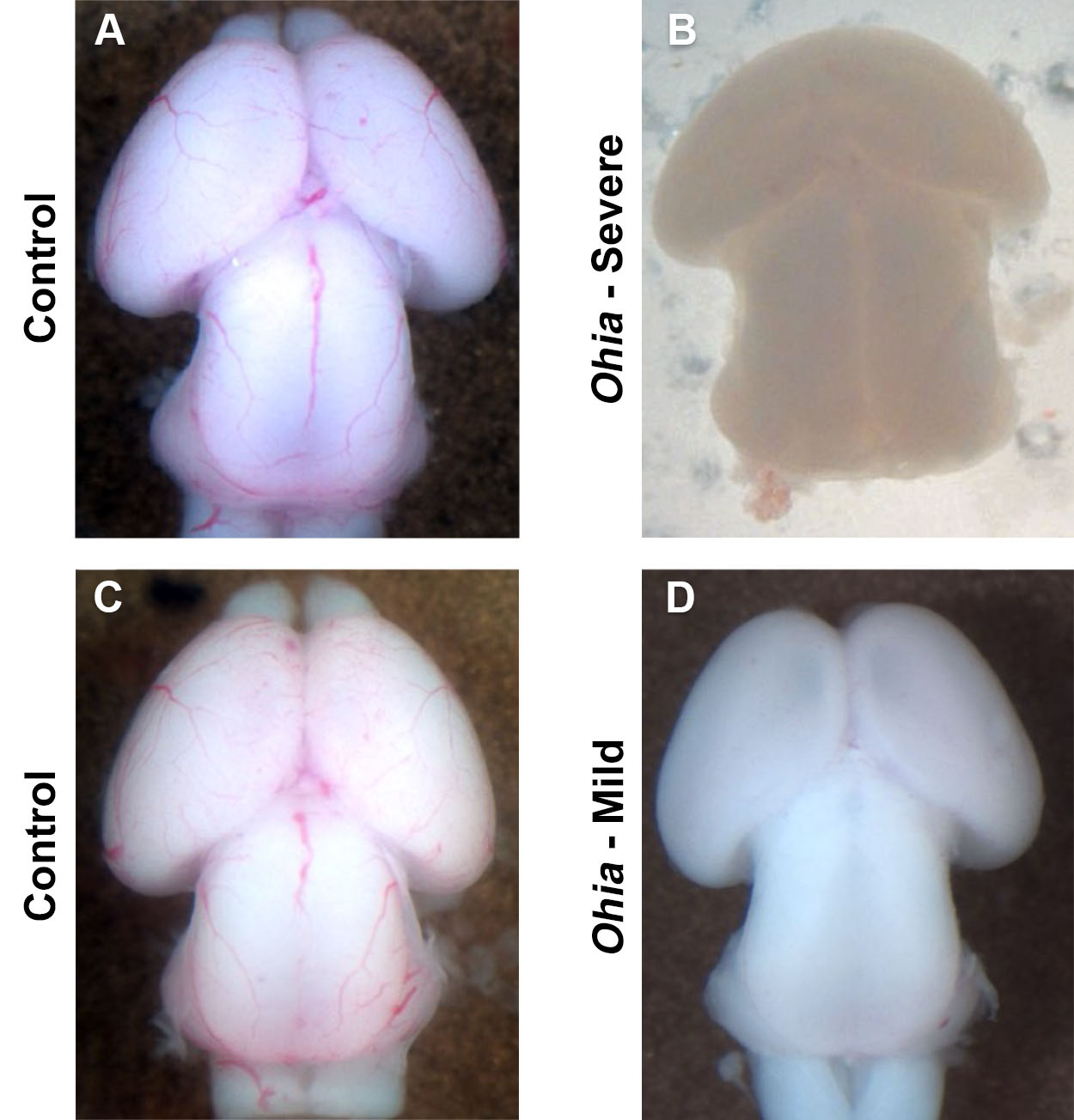

D

CA

**Figure S4. DNA methylation analysis of *Ohia* mouse brain in mutant mice with severe vs. mild brain phenotype.**

DNA methylation analysis was conducted on DNA obtained from brain tissue of *Ohia* mutant mice with severe brain phenotype that can include microcephaly and holoprosencephaly (B) as compared to control, and *Ohia* mutant mice with mild brain phenotype that are difficult to distinguish from (D) controls (C).

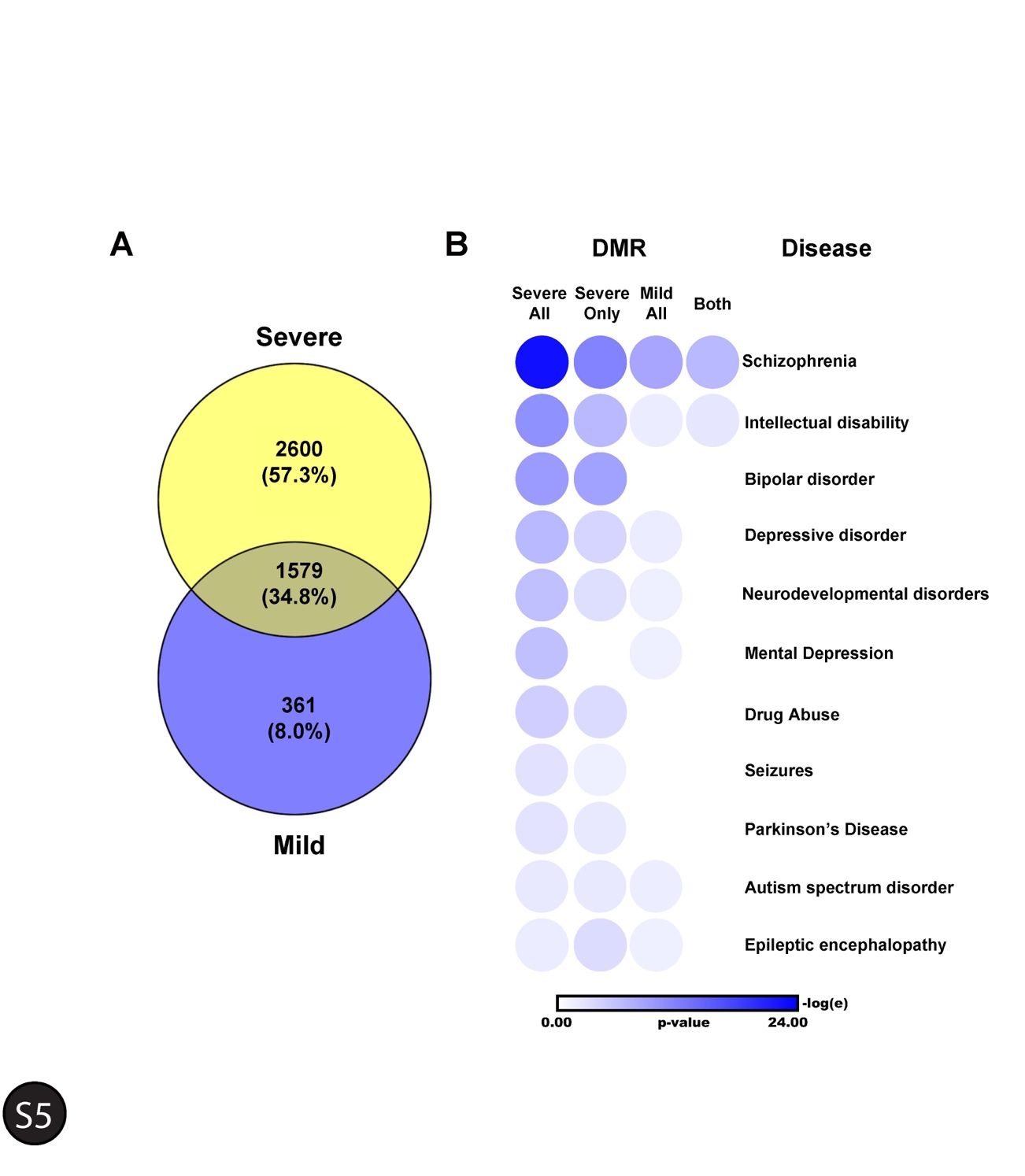

**Figure S5. Comparison of disease pathways recovered from analysis of differentially methylated regions in brain tissue of *Ohia* mutant mice with severe vs. mild brain defects.**

**A.** Brain tissue from *Ohia* mutant mice with severe and mild brain phenotypes were analyzed for DNA methylation changes relative to B6 wildtype control. Regions showing differential methylation, known as differentially methylated regions (DMR), were more abundant in the mutants with severe phenotype, with 2600 genes showing differential methylation exclusive to the severe phenotype. In contrast, the mutant with mild brain phenotype had DMRs associated with only 361 genes not observed in the mutants with the severe brain phenotype.

**B.** DMRs from *Ohia* mutant mice with severe phenotypes yielded many neurological diseases (Severe All). Similar analysis of *Ohia* mutant mice with mild brain phenotype (indicated as Mild All) recovered only 6 of the 10 neurological diseases seen with the severe phenotype. Analysis of DMRs exclusive to the severe phenotype (Severe Only) recovered 9 of the 10 disease terms identified with all DMRs from the severe phenotype. With similar analysis of DMRs exclusive to the mild phenotype (Mild only), none of the neurological disease terms were recovered (not shown). Examination of the 1579 DMRs that are shared between the mild and severe phenotype (Both) yielded only 2 of the 10 neurological disease pathways.

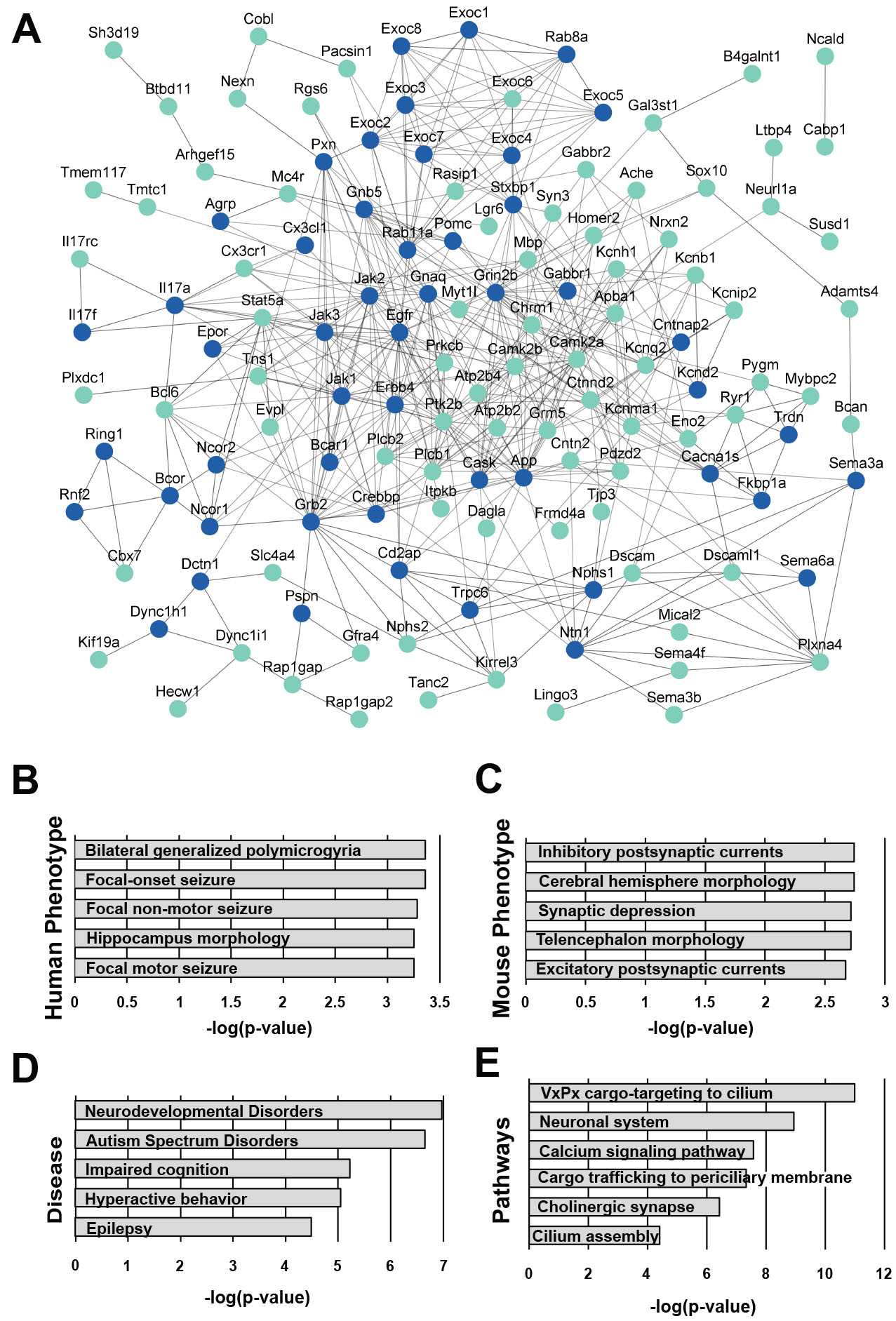

**Figure S6. Protein-Protein Interactome Network Analysis**

A protein-protein interactome was constructed for genes downregulated and differentially methylated (A). This network was then used in ToppGene analysis to identify human phenotypes (B), mouse phenotypes (C), diseases (D), and pathways (E) associated with this gene network.

**Figure S7. Behavioral Testing of *Pcdha9^m/m^* and EMX1-cre:*Sap130^f/-^* female mice.**

A. Morris water maze probe trial for female C57BL/6J WT Control and *Pcdha9^m/m^* animals.
B. Morris water maze visible platform trial for female C57BL/6J WT Control and *Pcdha9^m/m^* animals.
C. Contextual fear assessment in female C57BL/6J WT Control and *Pcdha9^m/m^* animals.
D. Time spent in each chamber over course of 10-minute trial in three chamber sociability test for female C57BL/6J WT Control and *Pcdha9^m/m^* animals.
E. Morris water maze probe trial for combined male and female control (EMX1-cre:*Sap130^f/+^* and *Sap130^f/-^*) and mutant EMX1-cre:*Sap130^f/-^* animals.
F. Morris water maze visible platform trial for for combined male and female control (EMX1-cre:*Sap130^f/+^* and *Sap130^f/-^*) and mutant EMX1-cre:*Sap130^f/-^* animals.
G. Contextual fear assessment in combined male and female control (EMX1-cre:*Sap130^f/+^* and *Sap130^f/-^*) and mutant EMX1-cre:*Sap130^f/-^* animals
H. Time spent in each chamber over course of 10-minute trial in three chamber sociability test for combined male and female control (EMX1-cre:*Sap130^f/+^* and *Sap130^f/-^*) and mutant EMX1-cre:*Sap130^f/-^* animals

**Figure S8. Behavioral Testing of *Pcdha9^m/m^* Male Mice.**

A. Morris water assessment over 5-day training period for male C57BL/6J WT Control and *Pcdha9^m/m^* animals.
B. Cued fear conditioning assessment of male C57BL/6J WT Control and *Pcdha9^m/m^* animals.
C. Sociability assessment of male C57BL/6J WT Control and *Pcdha9^m/m^* animals in the three chamber sociability test.
D. Morris water maze visible platform trial for male C57BL/6J WT Control and *Pcdha9^m/m^* animals.
E. Contextual fear assessment in male C57BL/6J WT Control and *Pcdha9^m/m^* animals.
F. Time spent in each chamber over course of 10-minute trial in three chamber sociability test for male C57BL/6J WT Control and *Pcdha9^m/m^* animals.
G. Morris water maze probe trial for combined male and female control (EMX1-cre:*Sap130^f/+^* and *Sap130^f/-^*) and mutant EMX1-cre:*Sap130^f/-^* animals.

**SUPPLEMENTAL METHODS**

**Mouse Husbandry**

*Ohia* and *Pcdha9* (c.2389_2399del; [p.Asp796Phefs∗]) mice generated by CRISPR gene editing were maintained in the C57BL/6J background. B6.129S2-Emx1tm1(cre)Krj/J mice with *Emx1-cre* driver were purchased from Jackson Laboratory (Strain #005628) and intercrossed with *Sap130^f/f^* and *Sap130^+/-^* mice generated previously^1^.

**Immunostaining and Confocal Microscopy**

*Ohia^m/m^* fetuses were obtained at E14.5, E15,5, and E16.5, the head was removed and drop fixed in 4% PFA overnight, and then processed for cryoembedding. Frozen sections were collected for immunostaining. Cryosections were stained with antibodies to Pax6 (BioLegend; 901301), Tbr1 (Abcam; ab31940), Tbr2 (EB Bioscience; 14-4875-80), Satb2 (Abcam; ab51502), Ctip2 (Abcam; ab18465), pH3 (Abcam; ab197502), Arl13b (Neuromab, N295B/66). TUNEL labeling was also performed with TUNEL assay kit from Roche (12156792910). Confocal imaging was conducted with the Leica SP8 confocal microscope and quantitatively analyzed with ImageJ. Figures were prepared with Adobe photoshop.

**Mouse Embryonic Fibroblast Analysis**

MEFs were isolated from E14.5-E15.5 embryos as previously described, with three independent MEF lines analyzed. Cells were plated on glass coverslips, fixed in 4% PFA, stained with antibodies to α-tubulin (Abcam; ab15246), and γ-tubulin (Sigma; T6557) and counterstained with DAPI (Thermo Fisher Scientific; D1306).

**Histological Reconstructions Using Episcopic Confocal Microscopy**

E14.5 or newborn mice were euthanized, the heads were removed, fixed in 4% paraformaldehyde, followed by embedding in paraffin for espicopic confocal microscopy (ECM) as previously described^1^. Briefly this entailed sectioning of the paraffin blocks using a Leica sledge microtome mounted with a LSI confocal macroscope that captures images of the block face at the microtome’s fixed photoposition. The images collected comprising a 2D serial image stack were sectioned 2D serial ECM image stacks were collected using a Leica sledge microtome are digitally resliced and also 3D reconstructed using the OsiriX Dicom viewer (<https://www.osirix-viewer.com>) to assess brain anatomical structures.

**RNA Sequencing**

Total RNA was isolated from whole brain tissue samples collected from 3 *Ohia^m/m^* animals with severe brain defects and 5 littermate control animals at E13.5-E14.5. Of the *Ohia^m/m^* animals used for this analysis, two showed HLHS heart defects while the third showed aortic defects. Additional analysis was conducted on 3 *Emx1-cre*:*Sap130^f/-^* and 3 littermate control *Sap130^f/-^* animals without Emx1-cre using the RNeasy plus mini kit (Qiagen). Libraries were constructed with a TruSeq RNA Sample Preparation Kit v2 (Illumina) and sequenced with an Illumina HiSeq 2000 platform (BGI Americas) with 100-bp paired-end reads. Reads were aligned to mm10 (NCBI build 38) with TopHat2 (v2.0.9)61, and gene-level counts were calculated with HTSeq-count (v0.5.4p5). Differential expression analyses were performed with edgeR, and mouse sex and litter were included as covariates in the analysis. Differentially expressed genes were recovered for *Ohia^m/m^* mutant tissue with false discovery rate (FDR) ≤0.05 (Benjamini–Hochberg) and no fold change cut off, and from *Emx1-cre*:*Sap130^f/-^* with FDR<0.1 and no fold change cut off. Pathway analysis was carried out using ToppFun from the ToppGene Suite^3^, Metascape^4^, and Ingenuity Pathway Analysis. Previously generated HLHS heart RNAseq results were used for comparison to the brain results identified in this study^1^. For the protein-protein interaction analysis, genes differentially expressed were recovered using FDR cut-off of 0.05. Genes with logFC>0 were considered up-regulated while logFC <0 were considered down-regulated.

**Chromatin Immunoprecipitation Sequencing**

Sap130 chromatin immunoprecipitation was performed as previously described^1^. E13.5 embryonic brains from wildtype C57BL/6J mice were used for analysis. Cross-linking was performed with 1% formaldehyde and stopped with glycine, and samples were sonicated to 100- to 300-bp fragments with a Covaris S2 instrument. Sonicated lysates were incubated with Sap130 antibody-bound Protein A magnetic beads (2 μg antibody per 20 μl beads). Sap130 chromatin immunoprecipitation was performed with rabbit anti-Sap130 antibody (A302-491A, Bethyl laboratories) and an iDeal ChIP–seq Kit for Transcription Factors (Diagenode). ChIP–seq libraries were generated with an NEBNext Ultra DNA Library Prep Kit Illumina (NEB), and sequencing was carried out on an Illumina HiSeq 4000 platform (BGI Americas). Reads were aligned to the mouse reference genome (mm10) with Bowtie1 (version 1.1.2), and potential Sap130 target regions were identified with MACS1.4.2 software.Reads were aligned to mm10 and Sap130 target regions were identified with MACS1.4.2. Motif enrichment analysis was performed using MEME suite version 5.4.1.

**Genome-wide DNA Methylation Analysis**

For methylation analysis, forebrain tissue samples were collected from 3 *Ohia^m/m^* with severe head phenotype, 3 *Ohia^m/m^* with mild head phenotype, and 3 wildtype C57BL/6J control mice at E15.5 and flash frozen in liquid nitrogen. DNA was isolated using the QIAamp DNA mini kit (Quiagen) and analyzed using the Illumina Infinium mouse methylation beadchip to assess DNA methylation (Illumina; 20041558). Bisulphite conversion of 250ng DNA was carried out using the EZ DNA Methylation™ Kit (Zymo Research Corp., CA), and bisulfite DNA samples were applied to the Infinium arrays and hybridized 16-24 hours at 48ºC. After further posthybridization processing, the beadchip was scanned using an Illumina iSCAN and the data analyzed using Genome Studio. Raw intensities were read into R using the R package minfi (v 1.36.0), and R methylation manifest and annotation packages to use with minfi were created using the Manifest and Annotation files provided by Illumina.

Samples with >1% of sites with detection p-value >0.01 were removed. Normal-exponential out-of-band (NOOB) normalization was performed. Poor quality probes with >=20% of samples with detection p-value >0.01 were filtered, as well as probes on sex chromosomes, and those that overlap SNPs were removed. Samples were pooled using the shared set of probes and final beta values were calculated. Probe-level differential methylation analyses was performed using the limma package, including sex as covariate in the design matrix. Probes with an FDR <=0.1 were considered significantly differentially methylated. Further, differentially methylated regions (DMRs) were identified using the R package DMRcate (v 2.4.1), with sex as a covariate, with an FDR of 0.05. DMR with maxdiff > 0 were considered hypermethylated regions and those with maxdiff <0 were considered hypomethylated regions

**Protein-Protein Interaction Analysis of Downregulated Genes Using STRING-db**

Gene expression with logFC>0 were considered up-regulated, while logFC <0 were considered down-regulated, and those overlapping with hypermethylated regions were considered hypermethylated and those overlapping with hypomethylated regions were considered hypomethylated genes. Genes were categorized based on differential DNA methylation and RNA expression, with the analysis focusing on the downregulated genes, yielding hypermethylated/down regulated genes and hypomethylated down regulated genes. These two gene sets were then used to retrieve interacting genes using the String-db database (STRING, https://string-db.org) to predict functional protein-protein interactions, and the top 50 interactors retrieved were then incorporated into an expanded network which was used for ToppGene pathway enrichment analysis.

**MRI Imaging**

For *in vivo* MRI, mice were anesthesized using inhaled isoflurane and then maintained with 1-2 % Isoflurane in oxygen rgwb transferred to the animal bed for imaging. MRI was carried out using a Bruker BioSpec 70/30 USR spectrometer (Bruker BioSpin MRI, Billerica, MA, USA) operating at 7-Tesla with quadrature radio-frequency volume coil with inner-diameter of 35 mm. Different brain areas, including the hippocampus, cerebral cortex, cerebellum, corpus callosum and forebrain, were manually defined by 2 blinded independent operators in ITK-SNAP, and volume for each brain region was calculated.

**Mouse Behavioral Phenotyping**

Mouse behavior was assessed using three training paradigms: the Morris water maze, fear conditioning, and sociability. Mice were group housed by sex during testing. For mice that underwent all three of these tests, testing was completed in order of least stressful to most stressful with sociability being conducted first followed by the Morris water maze, and then fear conditioning. On the day of testing, mice were transferred to the testing facility and allowed to habituate for at least one hour prior to test start. Testing apparatuses were cleaned with 70% ethanol between each animal tested. Testing was completed by operator blinded to genotype. For all tests, mice were weighed on the day of testing, and mice with loss of >10% of baseline weight between test days were excluded from analysis. Additionally, if mice appeared ill including little to no movement, piloerection, abnormal posture, or trouble breathing, they were excluded from analysis. This resulted in exclusion of 3 mice total.

**Morris water maze** The Morris water maze set-up consisted of a pool of 90 cm in diameter and 60 cm high filled with water (depth 28 cm, 20-22 degrees C) located in a 2.5 x 2.5 meter room with multiple extra-maze cues (black shapes on white wall) that remained constant throughout the experimental studies. A hidden platform 10 cm in diameter and 27 cm high (i.e.,1 cm below the water’s surface) was used to assess the mouse’s ability to learn spatial relations between distal cues and escape via this platform. Mice were given 120 sec per trial to find the hidden platform, and if the mouse failed to find the platform after 120 sec it was placed on the platform. Following each trial, mice remained on the platform for 30 sec before being moved to a heated incubator between trials. Mice were given 4 trials per day, starting at each of the 4 quadrants of the pool, and were tested over a period of 5 days. On Day 6 a probe test was completed, and testing was also done with a visible platform. The water maze pool and incubator were cleaned and disinfected daily. A video tracking system (AnyMaze) in conjunction with a digital camera mounted over the pool was used to record and quantitate the swimming behavior of the animals.

**Fear Conditioning** The fear conditioning apparatus consisted of a box (30.5x24.1x21 cm, Medical Associates) with an electric grid floor (1.6 cm spacing, 4.8 mm diameter rods). Animals initially underwent a training protocol in which they were placed in the box and after 120 sec, a 10 sec tone (75 dB, 32.8 kHz) was given, and during the last second of this tone, a 1 mA 1 sec long footshock was delivered through the grid floor. The tone and footshock co-terminated. The tone and foot shock were given sequentially for a total training period of 9 min, with a 60 sec intertrial interval. 24 hours after training, contextual fear conditioning was assessed by placing the animals in the box, and freezing was measured for 3 min. One hour after the assessment of contextual fear conditioning, cued fear conditioning was assessed by placing the animals in an altered chamber (the box was cleaned and scented with a novel odorant, a white noise of 75 dB is given, and a paper roll lined with print is placed within the box to hide the walls). The animal’s freezing behavior was measured for 3 min during which the tone (75 dB, 32.8 kHz) is delivered for the last minute, yielding the freezing behavior measurement with “cue”. No footshocks were delivered during this assessment. Following the cue the animal is observed for an additional minute to measure “post-cue” freezing behavior. Freezing behavior was quantified using a video-based analysis system (FreezeFrame, Coulburn Instruments) and verified by manual scoring.

**Sociability** Mouse social behavior was examined using a three-chambered social approach task. We used a video camera and EthovisionXT video tracking software (Noldus) to follow subject mice as they freely explored a three-chamber sociability cage. A test mouse was placed into the center compartment of the sociability cage, which is a box consisting of three compartments. In the other two compartments, we placed either a mouse with which the test mouse was unfamiliar, or an empty cage. At the beginning of the experiment the test mouse was placed into the middle chamber and habituated for 5 minutes. Following habituation, the novel mouse and empty cage were placed into either side of the three-chamber apparatus and the test mouse was able to freely explore each compartment for 10 minutes. The amount of time the test mouse spent with the empty cage versus with the unfamiliar mouse was quantified.

**Statistical Analysis**

Data were analyzed using GraphPad Prism 9 (GraphPad Software). For cell quantification independent samples T-test was used. For mouse behavioral analysis, data were analyzed by two-way, two-way repeated measures, or three-way analysis of variance (ANOVA). A p-value less than 0.05 was considered significant (p<0.05).

**Study Approval**

Mouse studies were conducted under an animal study protocol approved by the University of Pittsburgh Institutional Animal Care and Use Committee.

**SUPPLEMENTAL REFERENCES**

1.	Liu X, Yagi H, Saeed S, Bais AS, Gabriel GC, Chen Z, Peterson KA, Li Y, Schwartz MC and Reynolds WT. The complex genetics of hypoplastic left heart syndrome. *Nature genetics*. 2017;49:1152-1159.

2.	Xu X, Jin K, Bais AS, Zhu W, Yagi H, Feinstein TN, Nguyen PK, Criscione JD, Liu X, Beutner G, Karunakaran KB, Rao KS, He H, Adams P, Kuo CK, Kostka D, Pryhuber GS, Shiva S, Ganapathiraju MK, Porter GA, Lin J-HI, Aronow B and Lo CW. Uncompensated mitochondrial oxidative stress underlies heart failure in an iPSC-derived model of congenital heart disease. *Cell Stem Cell*. 2022.

3.	Chen J, Bardes EE, Aronow BJ and Jegga AG. ToppGene Suite for gene list enrichment analysis and candidate gene prioritization. *Nucleic acids research*. 2009;37:W305-W311.

4.	Zhou Y, Zhou B, Pache L, Chang M, Khodabakhshi AH, Tanaseichuk O, Benner C and Chanda SK. Metascape provides a biologist-oriented resource for the analysis of systems-level datasets. *Nature communications*. 2019;10:1-10.
